# Therapeutic poxviruses induce the secretion of immunostimulating and anti-tumoral extracellular vesicles

**DOI:** 10.1101/2025.09.19.677320

**Authors:** Lucas Walther, Vincent Mittelheisser, Marie-Christine Claudepierre, Louis Bochler, Magali Rompais, Jules Deforges, Nathalie Silvestre, Annabel Larnicol, Christine Carapito, Eric Quémeneur, Jacky G. Goetz, Karola Rittner, Vincent Hyenne

**Author notes:** These authors contributed equally to this work (Jacky G. Goetz, Karola Rittner, Vincent Hyenne).

## Abstract

Poxvirus-based vectors provide a versatile cancer immunotherapy platform, enabling the expression of immunostimulatory molecules and cancer-specific antigens. While infections with pathogenic viruses are well known to modulate extracellular vesicle (EV) biogenesis and function, the extent to which therapeutic poxviral vectors influence EV secretion by immune cells and thereby affect therapeutic efficacy remains underexplored. In this study, we showed that poxviruses, including the clinically relevant Modified Vaccinia Ankara (MVA), stimulate the secretion of small EVs (sEVs) containing viral proteins and immune-related signatures from peripheral blood mononuclear cells (PBMCs). Using an engineered MVA vector, we demonstrated the transfer of virus-encoded therapeutic payloads to sEVs, including the model ovalbumin (OVA)-derived peptide SIINFEKL presented by the class I major histocompatibility complex (MHC I) and the immune activators interleukin-12 (IL-12) and CD40 ligand (CD40L). Depending on the isolation method, these sEVs stimulated SIINFEKL-specific CD8⁺ T cells with varying efficiencies *in vitro*. Remarkably, intravenous injection of these sEVs into E.G7-OVA lymphoma–bearing mice reduced tumor growth to an extent comparable to the virus itself. Taken together, our findings indicate that EVs released from immune cells infected with engineered therapeutic poxviruses exert potent antitumor activity. These vesicles represent actionable mediators whose secretion and functionalization can be harnessed to improve viral vector–based immunotherapies, as well as being considered as therapeutic vectors in their own.

**Graphical Abstract:** 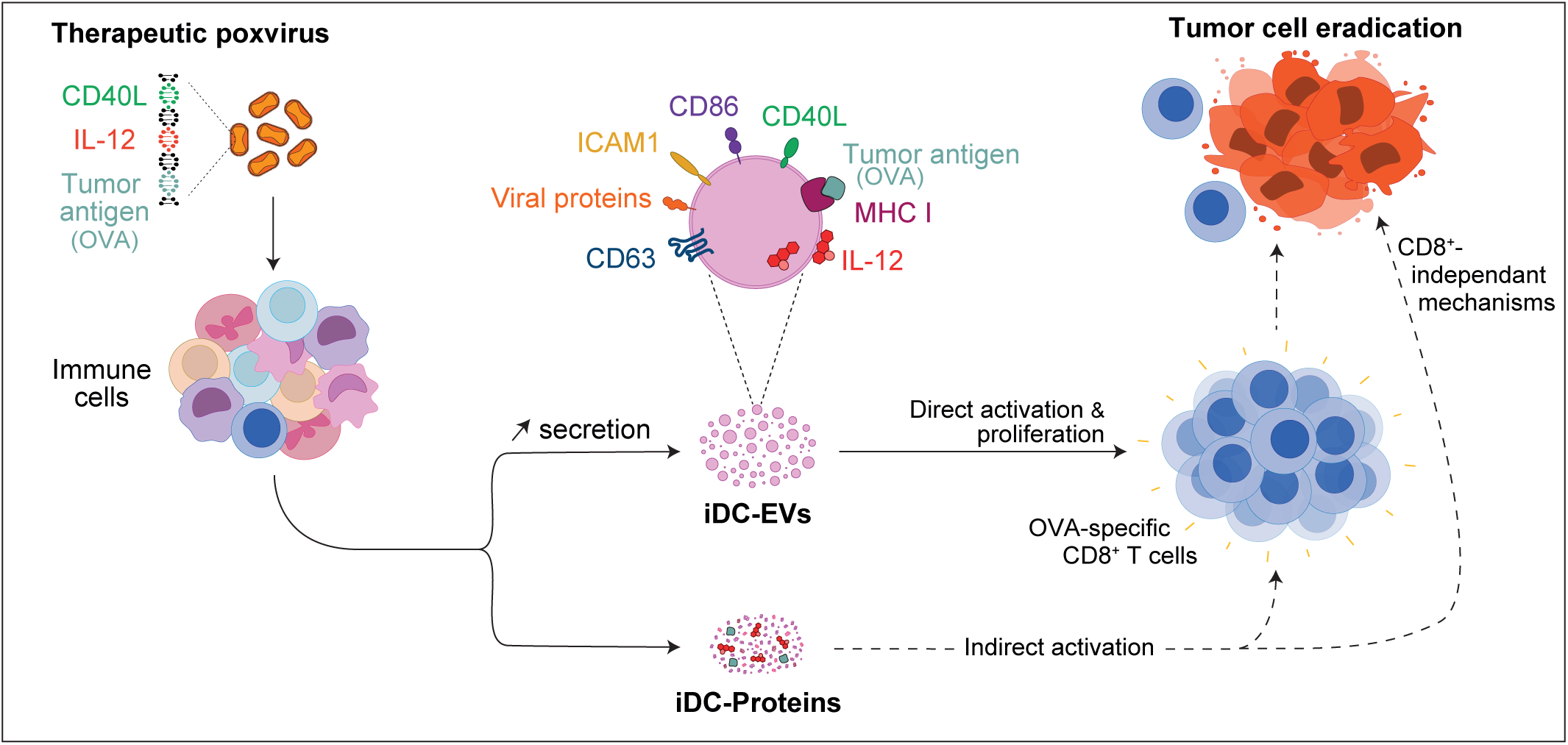

## Introduction

Virus-based vectors, including lentiviruses, adenoviruses and poxviruses (Bezeljak, 2022; Bendjama and Quemeneur, 2017), have emerged as a versatile and clinically validated platform for gene delivery, oncolytic virotherapy, and therapeutic vaccination in oncology. For instance, engineered poxviral vectors eradicate tumors by (i) enhancing tumor tropism; (ii) conferring tumor-selective replication (Foloppe *et al*., 2019); (iii) expressing tumor-associated antigens (TAAs) or (iv) delivering immunostimulatory molecules (Travieso *et al*., 2022). They also support therapeutic cancer vaccination, including personalized neoantigen constructs that elicit robust, patient-tailored immune responses (Seclì *et al*., 2023). Among them, the Modified Vaccinia Ankara (MVA) is an attenuated poxvirus strain that has been extensively investigated and demonstrated to be a safe, flexible, and immunogenic vector (Cottingham et Carroll, 2013). In preclinical studies, MVA vaccination was shown to induce the generation of tumor antigen-specific CD8^+^ T cells able to trigger a complete regression of tumors (Ramos *et al*., 2022). The MVA-based personalized vaccine TG4050 targeting neoantigens in head and neck cancer is currently under clinical evaluation (ClinicalTrials.gov identifier NCT04183166). Yet the full consequences of therapeutic virus infection, both on the infected immune cells and the subsequent events leading to adaptive immunity activation, are incompletely understood. A deeper understanding of how therapeutic viral infection remodels host cells and shapes systemic immunity will not only enable the development of safer and more effective treatments but also broaden our understanding of the mechanisms by which viruses modulate infection efficacy, together with the magnitude and quality of the elicited immune response. Notably, one broadly overlooked consequence of viral infection is the remodelling of extracellular vesicles (EVs) secretion and content, reflecting the complex, intertwined links between viruses and EV biogenesis pathways (Martin *et al*., 2024). EVs are heterogeneous particles delimited by a lipid bilayer (ranging from ≈30 nm to a few µm) and secreted by most cells via different pathways, from endosomes or from the plasma membrane (Van Niel *et al*., 2018). As mediators of intercellular communication in physiology and disease (e.g., inflammation, cancer, cardiovascular disorders), EVs transport lipids, proteins and nucleic acids (RNA, DNA), with cargo composition reflecting the state and identity of the donor cell (Kalluri et LeBleu, 2020). On a translational level, EVs are being used both as circulating disease biomarkers and developed as versatile delivery vehicles for bioactive therapeutics (Kumar *et al*., 2024), notably in cancer, with growing emphasis on engineering for improved drug delivery and immunomodulation (Yang *et al*., 2024; Mohamed *et al*., 2025). Since EVs play a major role in modulating immune responses (Buzas, 2023), notably by presenting antigens at their surface to activate T cells (Greening *et al*., 2023), they emerged as vaccination vectors. Viruses are ontogenetically, structurally and functionally closely related to EVs (Cortes-Galvez *et al*., 2023; Moulin *et al*., 2023) and exploit their secretion mechanisms to their advantage (Hassan *et al*., 2021; Chatterjee *et al*., 2024). For instance, the vaccinia virus (strain WR) hijacks the endosomal sorting complex required for transport (ESCRT)-mediated multivesicular body (MVB) pathway, known to be involved in cargo sorting and exosome secretion (Xu *et al*., 2023), for its egress from the cell (Huttunen *et al*., 2021). As a consequence, infected cells secrete a variety of particles forming a continuum, ranging from unaltered host-derived EVs to fully infectious viral particles, with intermediates such as virus-induced EVs and virus-like particles (Nolte-‘t Hoen *et al*., 2016). For instance, cells infected with an MVA encoding the immunostimulatory molecule CD40L release EVs displaying both the viral envelope protein B5 and CD40L on their surface (Spehner and Drillien, 2008).

Based on these observations, we wondered whether i) immunomodulatory therapeutic poxviruses could influence both the secretion and content of EVs secreted by infected immune cells and ii) such EVs may contribute functionally to the efficacy of anti-tumoral therapeutic vaccines. To test these hypotheses, we used three members of the *Poxviridae* family exploited for therapeutic purposes: the *Vaccinia Virus* (strain Copenhagen, VV_COP_) which is replication-competent; the Modified Vaccinia Ankara (MVA), an attenuated strain replication-deficient in most mammalian cells, and the Pseudocowpox virus (PCPV) that replicates efficiently in bovine cells. We first established a method to successfully separate EVs secreted by PBMCs, either infected with VV_COP_ or MVA, from closely related viral particles, based on size filtration and ultracentrifugation. Then, using this protocol, we characterized *in vitro* and *in vivo* the content and function of virus-free small EVs (sEVs) secreted by PBMCs or murine dendritic cells DC2.4 infected with surrogates MVA vectors used for preclinical studies, either encoding reporter gene eGFP (MVA-eGFP) or therapeutic payloads (^armed^MVA). We demonstrated that infection with the ^armed^MVA stimulates the release of sEVs loaded with MVA-encoded immunomodulatory payloads which prevent tumor progression *in vivo*. Our results suggest that EVs partially mediate the therapeutic effect of immunostimulatory poxviruses, paving the way for improved virus-based therapies by tuning EV-mediated functions or directly using EVs as therapeutic vehicles.

## Results

### Size filtration allows the separation of small EVs from infectious viral particles

Aiming at analysing the immunomodulatory potential of EVs secreted by cells infected with immunotherapeutic poxviruses, we first investigated which isolation method would reliably separate them from viral particles. This step is challenging, as poxviruses are dsDNA enveloped viruses, approximately 200–300 nm in diameter (McInnes *et al*., 2023), that share size, structural and biophysical characteristics with EVs (Nolte-‘t Hoen *et al*., 2016; Cortes-Galvez *et al*., 2023). In order to selectively isolate EVs from infectious viral particles, which is mandatory for their analysis and further use, we compared some of the most classical and robust EV isolation methods: ultracentrifugation (UC), immunocapture, density gradient, velocity gradient, size exclusion chromatography (SEC) and filtration (**Fig.1A and S1A-C**) (Welsh *et al*., 2024). PBMCs from healthy donors were infected with a typical representative of the *Poxviridae* family (of which other versions are currently tested in therapy, ClinicalTrials.gov identifier NCT04725331): the VV_COP_, encoding eGFP reporter protein (VV_COP_-eGFP). Cells were infected at a multiplicity of infection of 1 (MOI=1) over night and secreted EVs (and other co-isolated particles, including remaining or newly-secreted viral particles) were isolated, measured and quantified by nanoparticle tracking analysis (NTA). The presence of residual infectious viral particles was probed by titration of plaque forming units (PFU).

**Figure 1:**
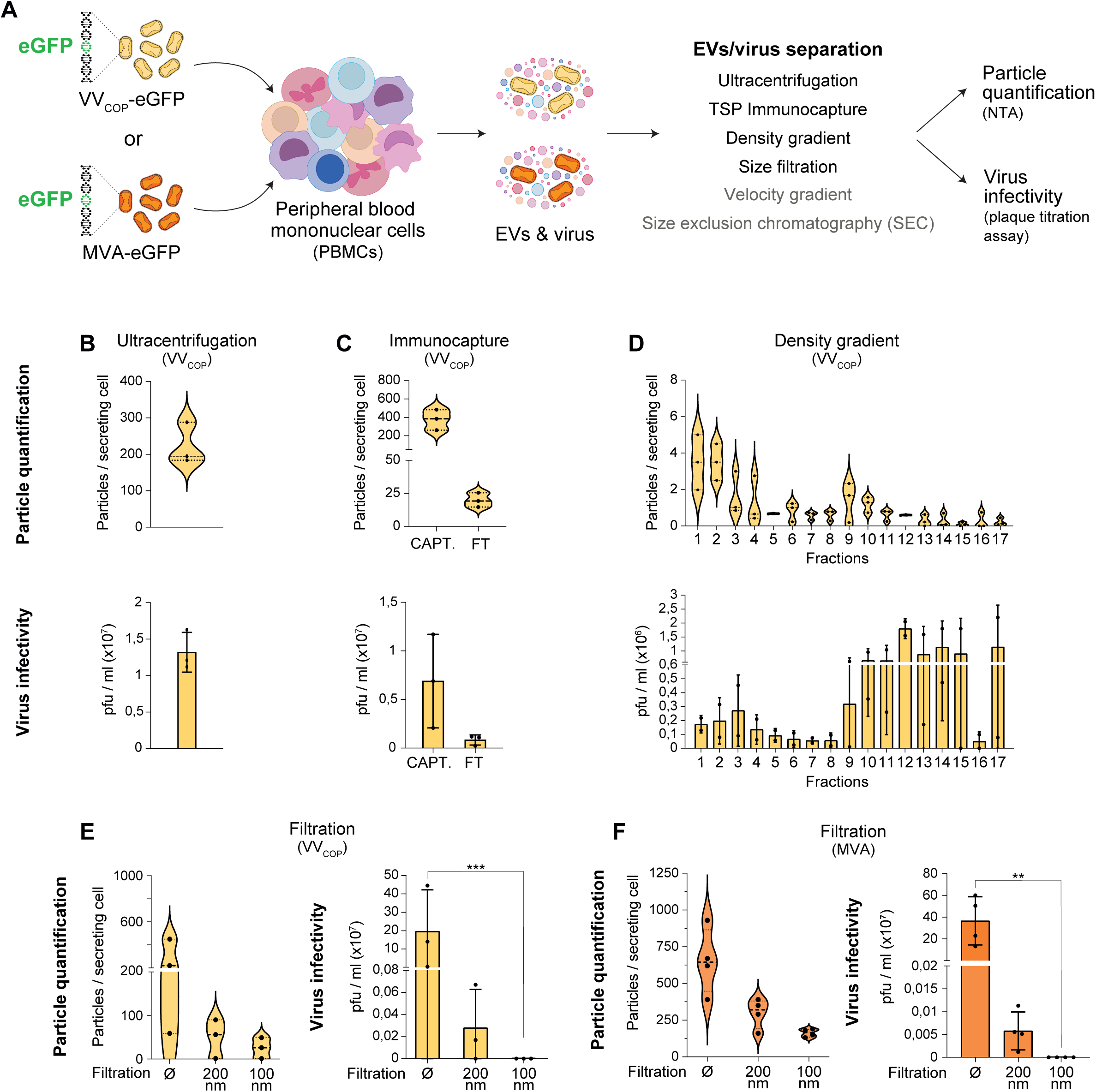
100nm filtration allows to efficiently separate small EVs from infectious poxvirus particles. **A)** Workflow for the comparison of six different EV isolation methods tested for their ability to separate small EVs from infectious poxvirus particles from the supernatant of human peripheral blood mononuclear cells (PBMCs) infected by VV_COP_ **(B-E)** or MVA **(F)** poxvirus at a multiplicity of infection of 1 (MOI=1). Particles were quantified by NTA and virus infectivity was assessed by titration of plaque forming units (PFU). **B-F)** Top: quantification of the number of isolated particles per secreting cell; Bottom: quantification of the virus infectivity. Supernatants from poxvirus-infected PBMCs were processed by ultracentrifugation **(B),** N=3 independent experiments; pan-tetraspanin immunocapture **(C)** (CAPT.: captured particles; FT: flow through), N=3 independent experiments; iodixanol density gradient **(D),** N=3 independent experiments; and filtration **(E-F)** at 200 nm (0.2µm) or 100 nm (0.1µm), N=3 independent experiments for VV_COP_ in **(E)**, Filtered 100nm p=0,019, Kruskal-Wallis; and N=4 independent experiments for MVA in (**F**), Filtered 100nm p=0,0028, Kruskal-Wallis. ** p<0,05; *** p<0,005.

We first observed that ultracentrifugation at 100,000g co-pelleted both EVs and infectious viral particles (**Fig. 1B**). In order to visualize EVs, we performed electron microscopy and confirmed that such method cannot separate EVs from brick-shaped viral particles that have overlapping size and density (**Fig. S1A**). We next tested a method relying on the immunocapture of tetraspanins (CD9, CD81 and CD63), which are classically present on the surface of EVs (Welsh *et al*., 2024). While this approach efficiently captured high amount of particles, it also retained most of the infectivity present in the samples (**Fig. 1C**). This may be explained by the fact that poxviruses, particularly in their enveloped form, known as extracellular enveloped virus (EEV), originate at least in part from tetraspanin-rich compartments, such as multivesicular bodies (Huttunen *et al*., 2021) and incorporate host cell components in their envelop (Vanderplasschen *et al*., 1998).

We then tested fractionation methods based on the structural properties of the particles, hoping to identify fractions containing EVs and deprived of infectious particles. To this end, we tried to separate EVs from viral particles based on their density, using an iodixanol density gradient. The most infectious fractions (fractions 10-15) were distinctly separated from the fractions enriched in particles (fractions 1-4 and fractions 8-10) (**Fig.1D**). Interestingly, viral infection is associated with the apparition of low-density particles (fractions 1-4) distinct from usual EV fractions (fractions 8-10) (**Fig.S2C)** which could correspond to previously described EVs-virus intermediates (Nolte-‘t Hoen *et al*., 2016). Yet, infectious viral particles were identified in each of the 17 fractions collected, thereby excluding density gradient centrifugation as an efficient method to separate EVs from viral particles. We next used velocity gradient, which was recently shown to segregate HIV virus from small vesicles (Martin-Jaular *et al*., 2021), and size-based fractionation using size-exclusion chromatography (**Fig.S1B-C**). These methods equally failed to separate infectious poxvirus particles from EVs, demonstrating that neither size-nor density-based methods fractionation can efficiently overcome the shared characteristics of poxvirus particles and EVs. Finally, we investigated whether a size cut-off filtration (*i.e.* positive enrichment of small sized particles) could efficiently separate small EVs (sEVs) from viral particles. To test this approach, we applied 200 nm or 100 nm filtration as an ultimate step following ultracentrifugation (**Fig. 1E-F**). While 200 nm filtration significantly reduced the viral load (around a thousand-fold), it failed to provide a purified particle preparation devoid of infectivity. In contrast, 100 nm filtration successfully yielded virus-free sEVs, with a diameter of approximately 130-150 nm as measured by NTA (**Fig. 1E**). Extending this approach to MVA, another widely used and well-characterised vaccinia virus strain, we showed it equally succeed in isolating sEVs free of infectious particles (**Fig. 1F**). However, such method remained inefficient to separate sEVs from the PCPV (*Parapoxvirus*), which appears more elongated than the brick-shaped VV_COP_ and MVA (*Orthopoxvirus*) as previously described (Gelderblom et Madeley, 2018) and might therefore not be fully retained by 100 nm pores (**Fig. S1D-E**). Altogether, this careful screen of particle isolation methods demonstrates that 100nm filtration coupled to ultracentrifugation is the only one efficiently segregating sEVs from any VV_COP_ and MVA infectious viral particles while allowing to collect sufficient amounts of virus-free sEVs. This protocol now allows for the first time to investigate the immunomodulatory potential specific to sEVs secreted by orthopoxvirus-infected cells.

### MVA infection enhances EVs secretion by PBMCs and alters their content

Building on a reliable method to study sEVs from infected cells, we then investigated the impact of poxvirus infection on EV secretion levels and content (**Fig. 2A**). In our subsequent work we focused on the attenuated *Orthopoxvirus* strain MVA since this virus maintains a strong immunogenicity profile and serves as the backbone for MVA-based vectors currently under clinical evaluation as therapeutic cancer vaccine. PBMCs from healthy donors were infected with MVA-eGFP, allowing to track virus-encoded payload and EVs deprived of infectious viral particles were isolated by UC combined with 100 nm filtration and quantified by NTA. We found that MVA infection leads to a five-fold increase in the number of sEVs compared to a non-infected condition (**Fig. 2B**). Such increase was observed independently of the isolation method (UC (**Fig. S2A**), density (**Fig. S2C**) or velocity gradient (**Fig. S2D**), and SEC (**Fig. S2E**)) and the poxviruses strain tested (**Fig.S2F**), suggesting that poxvirus-infected PBMCs significantly increase their EV secretion levels. We next wondered whether the infection had any impact their content. Mass spectrometry analyses of EVs from poxvirus-infected PBMCs revealed that MVA-eGFP infection markedly alters the protein composition of sEVs (**Fig.2 C-E, Fig.S3**). GO terms analysis of the differentially expressed proteins revealed an enrichment for terms associated with EVs, host-virus interactions and adaptative immunity (**Fig. S3A**). Around 50 PBMCs cellular proteins displayed a differential expression, most of them showing an increase post-infection (**Fig.2 C, D; Fig.S3B and Tables 1-3**). Among them, we found proteins related to viral response (SAMHD1, Rack1, Magt1), translation (RPL5, EIF3D, RPS9), vesicular trafficking (Rab7, VPS35) and actin polymerization (Espin, Arpc4, Profilin, Myo1A). In addition, we identified various viral proteins associated with EVs upon the different filtration methods. While their presence in the unfiltered samples could be explained by the co-isolation of viral particles (**Fig.2 D-E, Fig.S3C, Table 1**), their presence after the 100nm filtration reveals that some viral proteins are integrated at the surface or inside sEVs secreted after infection (**Fig.2 D-E, Table 3**). Among them, we identified core viral proteins, membrane and envelope proteins as well as proteins involved in viral replication and in immune modulation (**Fig.2E**). Notably, we also observed the presence of virus-encoded eGFP in sEVs after 100nm filtration (**Fig.2 D, E)**. Overall, we show that poxvirus (MVA) infection not only stimulates EV secretion in infected cells, but also significantly modifies their content, with upregulation of cellular proteins associated with viral infection and replication, as well as incorporation of various viral proteins. Most importantly, we demonstrated that exogenous payloads encoded by the virus (eGFP) can be efficiently transferred to sEVs secreted by infected cells, which opens the door to studying the transfer of immunomodulatory payloads to EVs, as a mean to leverage the therapeutic effect of the viral vector.

**Figure 2:**
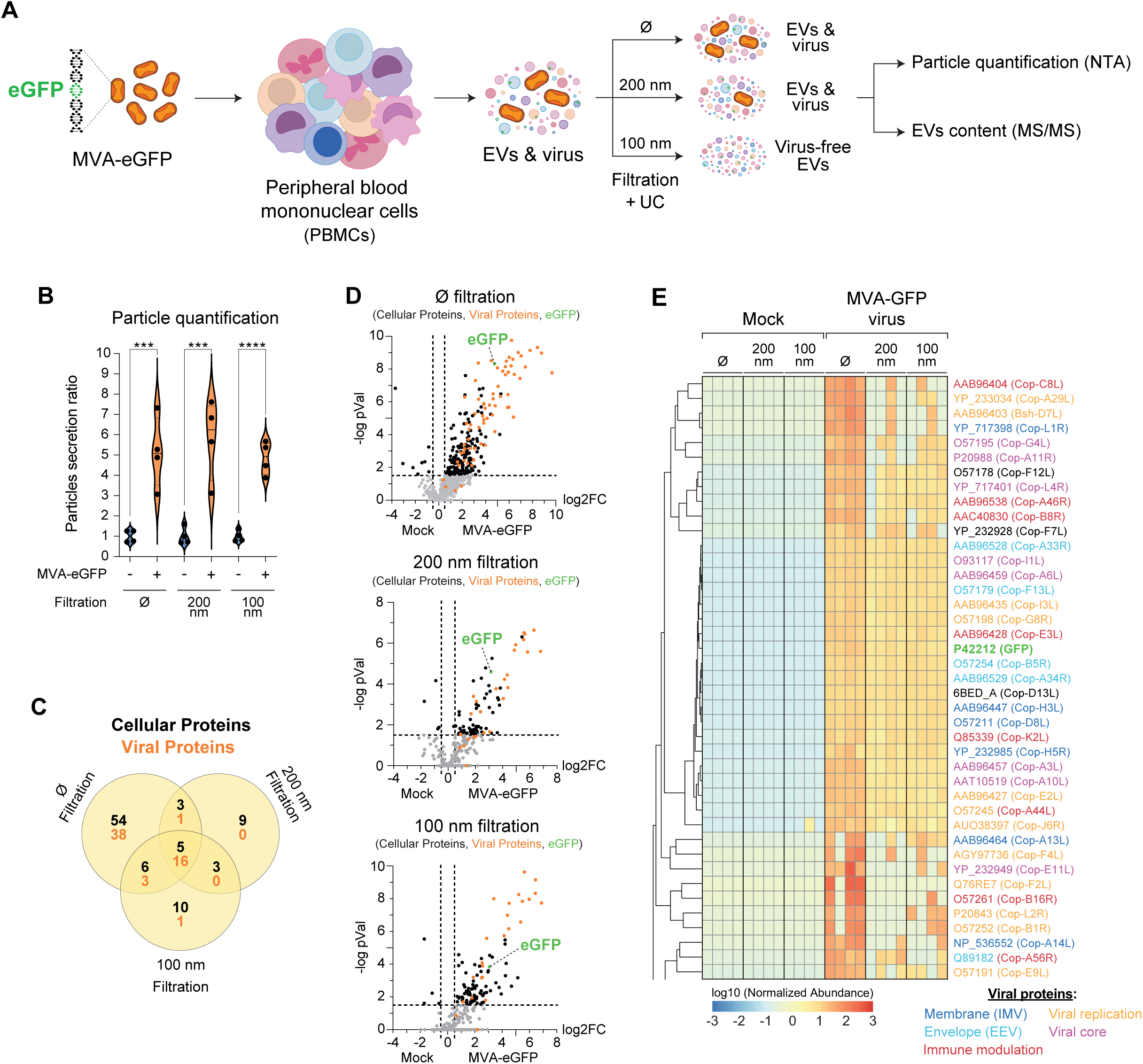
MVA infection increases the secretion of small EVs with distinct protein content. **A)** EVs secreted by human peripheral blood mononuclear cells (PBMC) infected by eGFP-MVA poxvirus were isolated by ultracentrifugation after 100nm or 200nm filtration, quantified by NTA **(B)** and their content was characterized by mass spectrometry **(C-E)**. **B)** Quantification by NTA of the number of particles per secreting cell in non-infected mock (blue) and MVA-infected (orange) conditions with and without filtration (normalized to mock). N=4 independent experiments. Unfiltered p=0,0034; Filtered 200nm p= 0,003; Filtered 100nm p=0,0001, Unpaired T-test. **C)** Venn diagram showing viral (orange) and cellular (black) proteins identified in ultracentrifugation pellet, with or without filtration (100nm or 200nm). **D)** Volcano plots showing viral (orange) and cellular (black) proteins identified in EVs isolated by ultracentrifugation, with or without filtration (100nm or 200nm) and differentially expressed between non-infected mock and MVA-infected conditions. Proteins with a p-value <0,01 and a log2FC>0,5 were considered. Virus-encoded eGFP is highlighted in green. **E)** Heat map showing the normalized abundance of viral proteins identified in ultracentrifugation pellet with or without filtration (100nm or 200nm) and color-coded by functional category. Label corresponds to MVA Uniprot ID (VVcop ortholog). Values are displayed as log10-transformed, normalized and centered by gene, with the corresponding color scale shown in the legend below. For proteins undetected by mass spectrometry in a given condition, the value was arbitrarily set to 0. Mass spectrometry was performed in quadruplicate. *** p<0,005; **** p<0,001.

### Dendritic cell–derived EVs carry therapeutic payloads encoded by an armed MVA

Prompted by the observation that virus-encoded eGFP can be transfered to sEVs, we next investigated whether therapeutic virus-encoded immunomodulatory payloads could be similarly transferred. In order to investigate payloads with proven immune-modulatory antitumor efficacy, we exploited the E.G7-OVA mouse lymphoma model to first compare the efficacy of various versions of therapeutic MVA vectors (**Fig.S4)**. Our results demonstrate that the highest anti-tumoral efficacy is obtained when several payloads were combined into one armed MVA: i) two epitopes of ovalbumin (OVA) as model antigens, including the widely used MHC class I-presented strong SIINFEKL peptide agonist to elicit OVA-specific responses, together with ii) the soluble murine cytokine IL-12, endogenously secreted by mature antigen-presenting cells (APC) to promote CD8^+^ T cells differentiation into effector cytotoxic T lymphocytes (Mirlekar and Pylayeva-Gupta, 2021), and iii) the murine CD40L, a membrane-bound immunostimulatory molecule which binds to CD40 to promote dendritic cells (DC) maturation, cytokines production, and prime CD8^+^ T cell into cytotoxic T cells (Elgueta *et al*., 2009). Besides its greater antitumor effect, we also selected this triple-armed vector for the complementary localization of its payloads: IL-12 is supposed to be secreted whereas CD40L and SIINFEKL presented by murine MHC-I (H-2Kᵇ) upon intracellular processing should be membrane bound, allowing us to track their association with sEVs. For testing the immunomodulatory potential of sEVs upon viral infection, we targeted professional antigen-presenting DC, as DC-derived EVs are known for their ability to present co-stimulatory molecules and antigen/MHC complexes on their surface that are able to trigger T cells response (Schioppa *et al*., 2024). More importantly, DC are well infected by MVA vector (Liu *et al*., 2008) and DC-derived EVs can elicit specific anti-tumor immune responses (Matsumoto *et al*., 2020). To explore this potential, we opted for the murine dendritic cell line DC2.4 (Liu *et al*., 2022), which was established from a C57BL/6 background and thus compatible with our E.G7-OVA model. After infection with either the triple-armed MVA (^armed^MVA) or an empty MVA (^empty^MVA), sEVs were isolated from infected DC (iDC) by 100 nm filtration followed by UC, as previously described, yielding ^armed^iDC-EVs-UC and ^empty^iDC-EVs-UC, respectively. sEVs from non-infected cells processed identically served as controls (DC-EVs-UC). In addition, we replaced UC with SEC after the 100nm filtration, to further refine the separation (Sidhom *et al*., 2020; Soares *et al*., 2023) of sEVs (^armed^iDC-EVs-SEC) from soluble proteins (^armed^iDC-Prots-SEC) (**Fig. 3A**). To characterise the surface composition of these various EVs we used an ELISA-like chemiluminescence immunoassay. Looking at markers of cellular origins, we observed reduced levels of the tetraspanin CD63, closely associated with endosomal-derived vesicles (Pols and Klumperman, 2009), at the surface of sEVs from infected DC2.4, independently of whether the virus was armed or not and independently of the EV isolation method (**Fig. 3B**). We also analysed two immunomodulatory proteins whose presence at the surface of DC promotes T cells activation: ICAM1 and CD86. ICAM1 is known for its important role in immune synapse strengthening by binding to LFA-1 (Bromley and Dustin, 2002; Sapoznikov *et al*., 2023; Zhou *et al*., 2025), allowing the stabilisation of the interaction between DC-derived EVs and T cells (Nolte-‘t Hoen *et al*., 2009) and it has been shown that its addition to the surface of engineered-EVs enhances mouse CD4^+^ and CD8^+^ T cell activation (Segura *et al*., 2005; Lyu *et al*., 2025). CD86 is a key co-stimulatory molecule involved in T cell activation through its interaction with CD28 (Thomas *et al*., 2007). EV surface expression levels were significantly increased for ICAM1 (**Fig.3C**) and a tendency was observed for CD86 (**Fig.3D**) upon MVA infection. Moreover, these three membrane-bound molecules were barely detected in the soluble protein-enriched fraction (^armed^iDC-Prots-SEC), validating the SEC efficacy. These results confirm that MVA infection induces a shift in sEVs identity, as initially observed with PBMC-derived EVs by mass spectrometry. We next assessed whether virus-encoded payloads would efficiently be transferred to the surface of EVs and observed that membrane-bound CD40L was exclusively detected on sEVs from DC2.4 cells infected with the ^armed^MVA (**Fig. 3E**). We subsequently demonstrated the presentation of the SIINFEKL peptide by mouse MHC class I (H-2K^b^) on the surface of sEVs derived from DC2.4 infected with the ^armed^MVA (**Fig. 3F**), using an antibody that detects only the SIINFEKL epitope when presented by mouse MHC class I. This is likely to support both direct and indirect antigen presentation to T cells, as well as cross-presentation to DC (Buzas, 2023). Finally, we quantified the murine IL-12 and found it in different proportion after infection with the ^armed^MVA. As expected, with approximately 100 times more IL-12, most of this soluble cytokine was recovered in the protein-enriched fraction (^armed^iDC-Prots-SEC), with a small amount still present in the sEVs fraction (^armed^iDC-EVs-SEC) following SEC isolation. Although some co-isolation can be observed with UC-isolated sEVs (^armed^iDC-EVs-UC), the majority was likely discarded in the supernatant during UC-based isolation (**Fig. 3G**). Taken together, our findings demonstrate that infection of DC with an armed MVA profoundly alters the molecular composition of their secreted EVs, resulting in (i) modified cargo content, (ii) enrichment of their surface with immunomodulatory cellular molecules such as ICAM-1, and (iii) incorporation of virus-encoded payloads including CD40L, MHC-I/SIINFEKL complexes, and small amount of IL-12. In conclusion, we demonstrated that the infection of mouse DC with an ^armed^MVA generates EVs with immunomodulatory potential likely to shape an anti-tumoral immune response.

**Figure 3:**
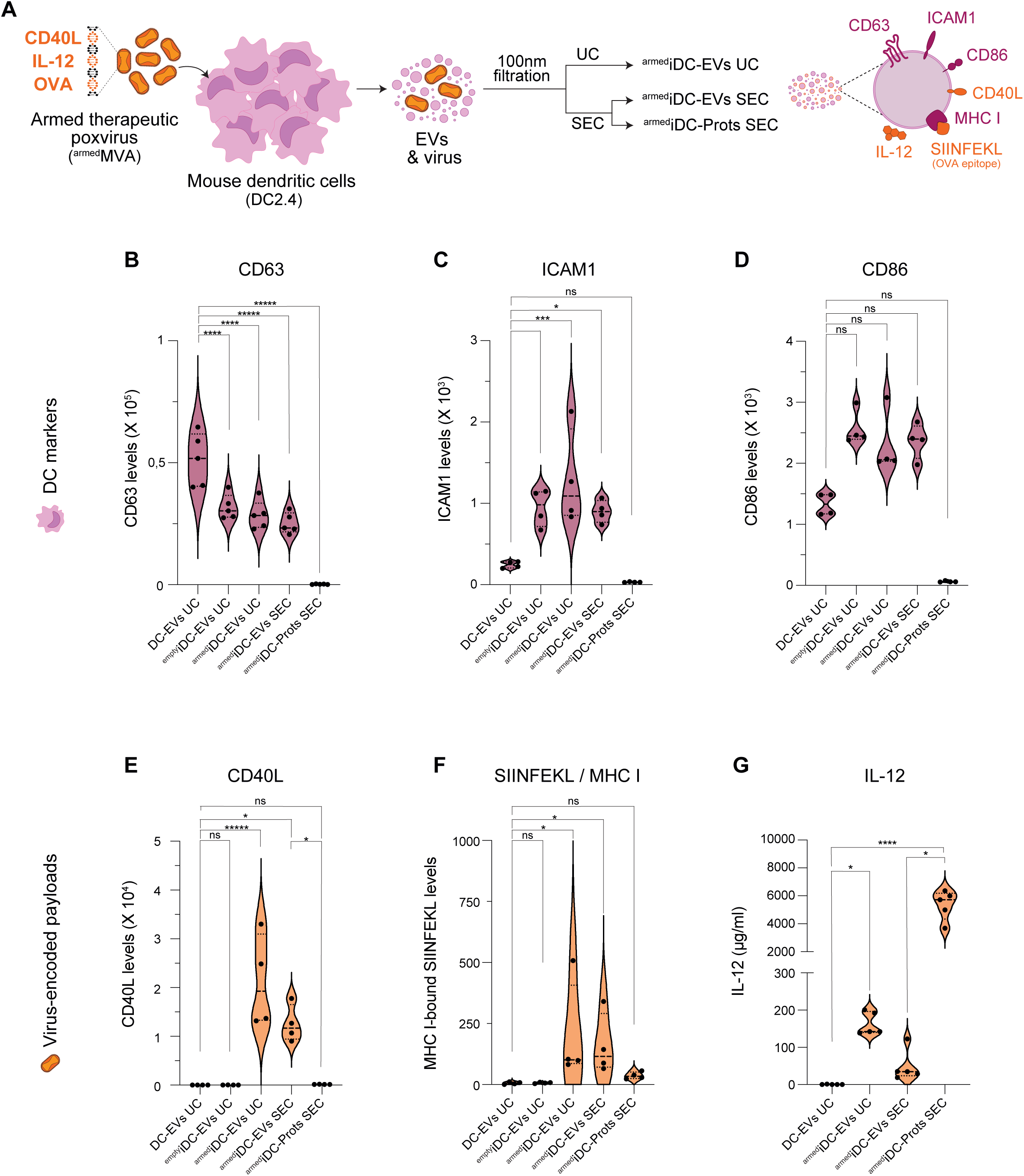
Virus-encoded therapeutic payloads are retrieved in small EVs. **A)** Workflow for assessing the presence of cellular proteins (CD63, ICAM1 and CD86) and virus-encoded therapeutic payloads (IL-12, CD40L and OVA peptide SIINFEKL presented on MHCI) on EVs secreted by mouse dendritic cells (DC2.4) and isolated by ultracentrifugation (UC) or size exclusion chromatography (SEC; EVs-rich fractions (EVs SEC, fractions 1-4) and soluble proteins-rich fractions (Prots SEC, fractions 5-10)) following 100nm filtration. Payloads were quantified by electrochemiluminescence immunoassay **(B-G)** on EVs isolated from non-infected cells (mock) or from cells infected with empty or armed MVA virus (encoding IL12, CD40L and OVA peptide SIINFEKL payloads). **B)** Quantification of CD63 on EVs. EVs mock Vs EVs empty virus p=0,0007; EVs mock Vs EVs armed virus UC p= 0,0001; EVs mock Vs EVs armed virus SEC p< 0,0001; EVs mock Vs Soluble proteins armed virus SEC p< 0,0001; One-way Anova; N=5 independent experiments. **C)** Quantification of ICAM1 on EVs. EVs mock Vs EVs empty virus: p=0,0276; EVs mock Vs EVs armed virus UC p= 0,0012; EVs mock Vs EVs armed virus SEC p= 0,0425; EVs mock Vs Soluble proteins armed virus SEC p= 0,83; One-way Anova; N=4 independent experiments. **D)** Quantification of CD86 on EVs. EVs mock Vs EVs empty virus: p=0,19; EVs mock Vs EVs armed virus UC p>0,99; EVs mock Vs EVs armed virus SEC p= 0,73; EVs mock Vs Soluble proteins armed virus SEC p= 0,99; Kruskal-Wallis; N=4 independent experiments. **E)** Quantification of CD40L on EVs. EVs mock Vs EVs empty virus: p>0,99; EVs mock Vs EVs armed virus UC p<0,0001; EVs mock Vs EVs armed virus SEC p= 0,011; EVs mock Vs Soluble proteins armed virus SEC p>0,99; EVs armed virus SEC Vs Soluble proteins armed virus SEC p=0,0125; One-way Anova; N=4 independent experiments. **F)** Quantification of MHCI bound OVA peptide (SIINFEKL). EVs mock Vs EVs empty virus: p>0,99; EVs mock Vs EVs armed virus UC p=0,023; EVs mock Vs EVs armed virus SEC p= 0,0341; EVs mock Vs Soluble proteins armed virus SEC p>0,99; Kruskal-Wallis; N=4 independent experiments. **E)** Quantification of IL-12. EVs mock Vs EVs empty virus p=0,045; EVs mock Vs EVs armed virus UC p>0,99; EVs mock Vs Soluble proteins armed virus SEC p=0,0004; EVs armed virus SEC Vs Soluble proteins armed virus SEC p=0,045; Kruskal-Wallis test. N=5 independent experiments. * p<0,05; ** p<0,01; *** p<0,005; **** p<0,001; ***** p<0,0001; ns: non-significant.

### sEVs secreted by dendritic cells infected with an armed MVA activate and expand CD8⁺ T cells

To assess this immunomodulatory potential, we next investigated whether sEVs secreted by ^armed^MVA-infected DC could directly stimulate SIINFEKL-specific CD8⁺ T cells *in vitro*. To this end, we isolated naïve CD8^+^ T cells from the spleens of OT-I transgenic mice, which express a TCR specific for the ovalbumin-derived peptide SIINFEKL presented by the murine MHC class I molecule H-2Kᵇ. These naïve CD8^+^ T cells were stimulated for 3 days with sEVs or the protein-enriched fraction secreted by ^armed^MVA-infected DC. We measured CD8^+^ T cell activation and proliferation by flow cytometry (**Fig. 4A**) and observed that ^armed^iDC-EVs UC induced a strong CD8⁺ T cell activation, as probed by CD44 expression, to levels comparable to the positive control stimulated with anti-CD3/CD28 antibodies (**Fig. 4B**). By contrast, sEVs from non-infected DC (DC-EVs UC) or from cells infected with an empty MVA (^empty^iDC-EVs-UC) failed to induce the activation of CD8^+^ T cells (**Fig. 4B**), while ^armed^iDC-EVs-SEC induced a moderate CD8^+^ T cells activation and the protein-enriched fractions (^armed^iDC-Prots-SEC), almost none. Further analysis of T cells proliferation revealed that ^armed^iDC-EVs-UC induced CD8^+^ T cells proliferation up to six cell generations (G0 to G6), with approximately twice as many cells in generations G5–G6, rising from 9,77% in the positive control to 18,18%. Consistent with their lack of activation, CD8^+^ T cells stimulated with either DC-EVs-UC or ^empty^iDC-EVs-UC mostly remained in generation G0, similar to the unstimulated control. On the other hand, ^armed^iDC-EVs-SEC and ^armed^iDC-Prots-SEC had weaker effects and show highly heterogeneous CD8^+^ T cell proliferation patterns (**Fig. 4C** and **Fig. S5A**). We further demonstrate that efficient CD8^+^ T cell activation relies on antigen presentation by sEVs, since MHC I/SIINFEKL blocking antibody (previously described by Blair *et al*., 2011) reduced CD8^+^ T cell proliferation in a dose-dependent manner (**Fig. S5B**). Altogether, our results demonstrate that sEVs isolated from DC2.4 infected with an armed MVA are able to activate specific CD8⁺ T cells and induce their proliferation *in vitro*, suggesting that they could also trigger a potent T-cell mediated antitumoral response *in vivo*.

**Figure 4:**
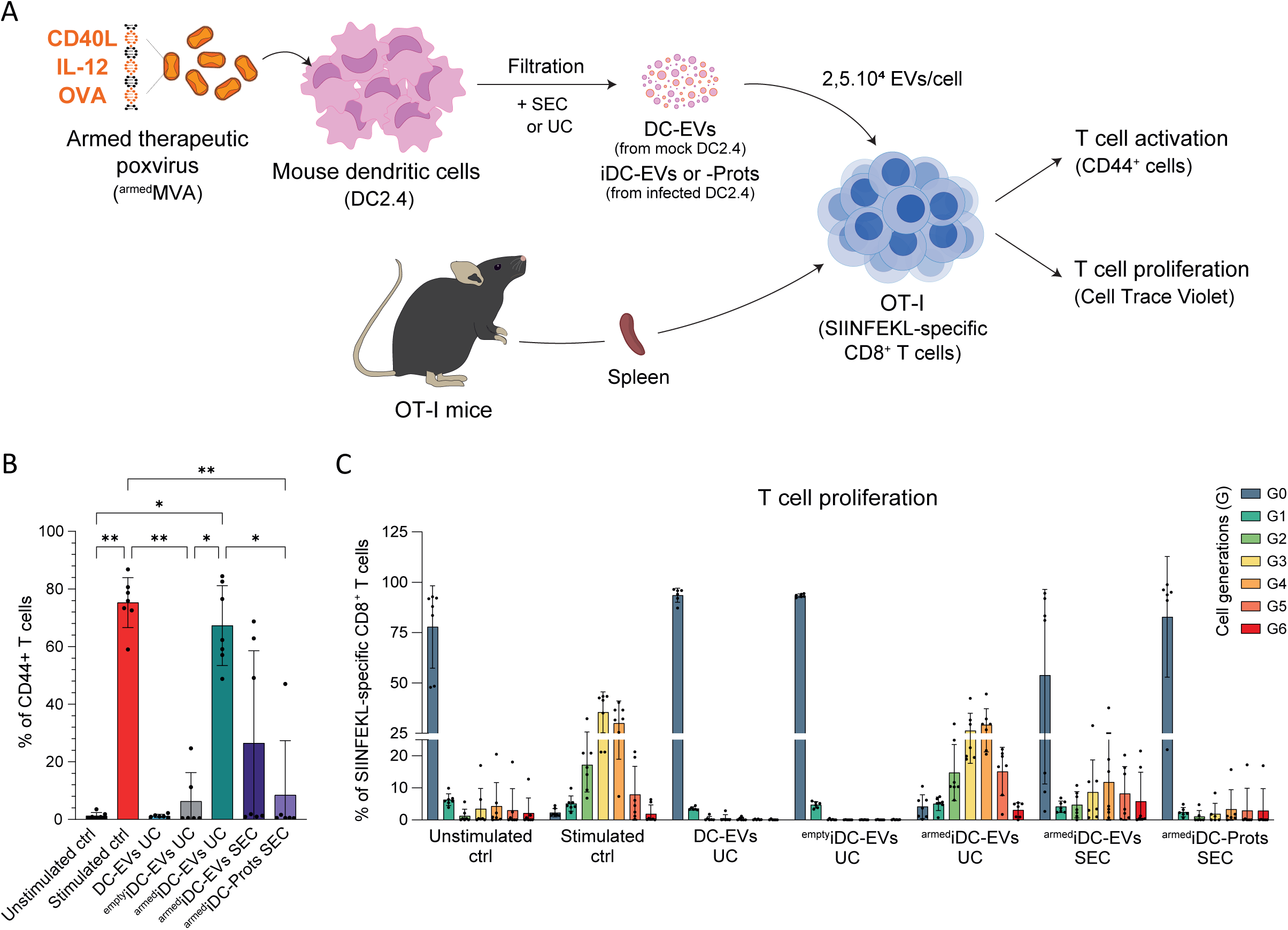
EVs isolated from therapeutic virus-infected dendritic cells activate and stimulate anti-tumoral cytotoxic T cells *in vitro*. A) Workflow for testing the ability of EVs to activate OVA-specific CD8^+^ T cells *in vitro* and stimulate their proliferation. EVs were isolated by ultracentrifugation (UC) or size exclusion chromatography (SEC) following 100 nm filtration, from non-infected murine dendritic cells (DC-EVs UC), or infected with an empty MVA virus (^empty^iDC-EVs UC), or with an armed MVA virus (encoding IL-12, CD40L and OVA peptide SIINFEKL payloads) yielding ^armed^iDC-EVs UC for EVs isolated by UC, ^armed^iDC-EVs SEC for EVs isolated by SEC and ^armed^iDC-Prots SEC for the soluble fraction isolated by SEC. PBS was used as a negative control (Unstimulated ctrl) and anti-CD3/CD28 antibodies as a positive control to activate T cells (Stimulated ctrl). **B)** Levels of T cells activation marker CD44 quantified by cytometry. Unstimulated ctrl Vs Stimulated ctrl p=0,0056; Stimulated ctrl Vs iDC-EVs UC (empty virus) p=0,0033; Unstimulated ctrl Vs iDC-EVs UC (armed virus) p=0,0245; Stimulated ctrl Vs iDC-Prots SEC (armed virus) p=0,0075; iDC-EVs UC (empty virus) Vs iDC-EVs UC (armed virus) p=0,0143; iDC-EVs UC (armed virus) Vs iDC-Prots SEC (armed virus) p=0,0303; Kruskal-Wallis. No significant difference in the absence of stars. n=1 to 2 replicates per group, in N=4 independent experiments. * p<0,05; ** p<0,01. **C)** Quantification of the number of T cells per generation (from G0 to G6) following stimulation with positive control or with EVs. n=1 to 2 replicates per group, in N=4 independent experiments.

### sEVs released by dendritic cells infected with an armed MVA prevent tumor growth

To assess the therapeutic potential of these sEVs secreted by ^armed^MVA-infected DC, we injected intravenously mice bearing E.G7-OVA lymphoma with sEVs or the protein-enriched fraction at days D7 and D14 post-tumor inoculation (**Fig. 5A**). PBS and the ^armed^MVA virus were used as controls. We observed that both, UC- and SEC-isolated sEVs from ^armed^MVA-infected DC, significantly reduced tumor growth, to levels obtained when mice were injected with the ^armed^MVA virus itself (**Fig.5B-C**). Notably, tumor growth patterns were very similar when mice were treated with ^armed^iDC-EVs-UC or the ^armed^MVA (**Fig. 5B**). Accordingly, survival of mice at D21 was significantly improved when treated with ^armed^iDC-EVs UC, compared to other treatments (**Fig. 5B**). While similar anti-tumoral efficacy was obtained with the protein-enriched fraction (^armed^iDC-Prots-SEC) obtained from ^armed^MVA-infected DC, sEVs from non-infected DC (DC-EVs-UC) had no effect on tumor growth. Measures at D21 revealed that sEVs or protein-enriched fraction from ^armed^MVA-infected DC could reduce tumor growth to levels obtained with the ^armed^MVA only (**Fig.5C**). To understand the underlying mechanism, we isolated and quantified at D21 the number of OVA-specific T cells in the peripheral blood of mice bearing an E.G7-OVA mouse lymphoma. While we detected SIINFEKL-specific CD8⁺ T cells in all groups treated with sEVs or protein-enriched fraction isolated from ^armed^MVA-infected DC (**Fig. S5C),** these were 10 to 50 times less abundant than in the ^armed^MVA virus-treated group. Altogether, our data demonstrate that the secretome (EV-bound and soluble) of DC infected by a fully armed MVA is as effective as the armed MVA alone in impairing the progression of the E.G7-OVA mouse lymphoma model, likely via controlling immunosurveillance.

**Figure 5:**
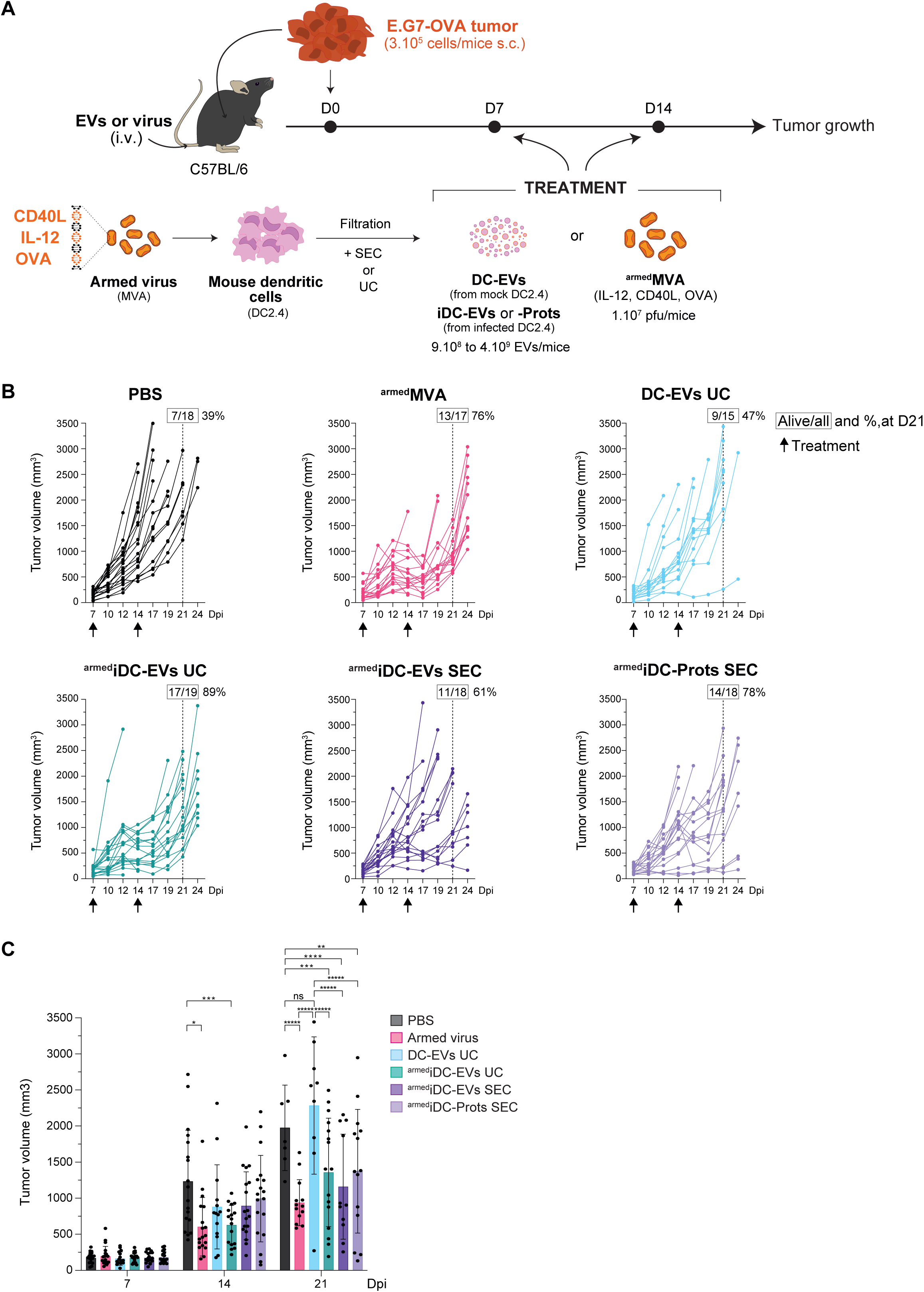
EVs isolated from therapeutic virus-infected dendritic cells inhibit tumor progression *in vivo*. **A)** Workflow for assessing the anti-tumoral potential of EVs secreted by mouse dendritic cells (DC2.4) isolated by ultracentrifugation (UC) or size exclusion chromatography (SEC; EVs-rich fractions (EVs SEC, fractions 1-4) and soluble proteins-rich fractions (Prots SEC, fractions 5-10)) following 100 nm filtration. EVs were isolated from non-infected cells (mock) or from cells infected with empty or armed MVA virus (encoding IL-12, CD40L and OVA peptide SIINFEKL payloads). EVs were injected intravenously 7- and 14-days post E.G7-OVA tumor cell injection and tumor growth was monitored for 24 days. **B)** Tumor growth curves of individual mice treated with PBS (black), with triple armed MVA virus (pink), with EVs isolated by ultracentrifugation from non-infected cells (mock, light blue) or from cells infected with armed MVA virus (green) or with EVs isolated by size exclusion chromatography (SEC), EVs rich fraction (iDC-EVs SEC, deep purple) or soluble proteins rich fraction (iDC-Prots SEC). Arrows indicate the injection days. Number and percentage of living mice at day 21 are indicated in boxes. One curve represents one mouse, n= 15 to 19 mice per group, in N= 4 independent experiments. **C)** Tumor volume at day 7, 14 and 21 post-tumor cell injection. At day 14, PBS Vs Armed virus p=0,0017; PBS Vs iDC-EVs UC (armed virus) p=0,019. At day 21, PBS Vs Armed virus p<0,0001; PBS Vs iDC-EVs UC (armed virus) p=0,0041; PBS Vs iDC-EVs SEC (armed virus) p=0,0005; PBS Vs iDC-Prots SEC (armed virus) p=0,0053; Armed virus Vs DC-EVs UC (mock) p<0,0001; iDC-EVs UC (armed virus) Vs DC-EVs UC (mock) p<0,0001; iDC-EVs SEC (armed virus) Vs DC-EVs UC (mock) p<0,0001 and iDC-Prots SEC (armed virus) Vs DC-EVs UC (mock) p<0,0001. Two-way Anova test. No significant difference with PBS in the absence of stars. * p<0,05; ** p<0,01; *** p<0,005; **** p<0,001; ***** p<0,0001; ns: non-significant.

## Discussion

Therapeutic vaccination is increasingly recognized as a new modality of immunotherapy, with accumulating preclinical and clinical evidence supporting its efficacy in inducing robust antitumor immune responses (Pail *et al*., 2025). Yet, the underlying molecular and cellular mechanisms remain incompletely characterised. In this study, we demonstrate that poxvirus-based vaccination induces the secretion of anti-tumoral sEVs by infected immune cells. Our *in vitro* and *in vivo* analyses indicate that these sEVs may constitute a key component of the vaccine-induced antitumor response, acting as mediators of antigen presentation and immune modulators, and may themselves hold promising potential as standalone therapeutic agents.

The similarities between EVs and viruses represents a significant challenge in the development of a universal separation method, due to their overlapping physical and biochemical properties, including size, density, membrane composition, and biogenesis pathways as well as the structural diversity among viral species (Nolte-‘t Hoen *et al*., 2016; Minh and Kamen, 2021). For these reasons, some of the methods approved with other viruses, such as density gradient for hCMV (Bergamelli *et al*., 2021) or velocity gradient for HIV (Martin-Jaular *et al*., 2021) have not proven effective for poxvirus in our study. By contrast, combining 100 nm filtration with UC or SEC effectively isolates sEVs from infectious MVA or VV_COP_ particles. While this approach allowed us to unambiguously demonstrate that small EVs released post-infection contain virus-encoded payloads and bear anti-tumoral immunomodulatory functions, it is also possible that bigger EVs possess similar properties. This method could be easily applied to the study of sEVs secreted by cells infected with other viruses exceeding 100 nm in size, such as the Epstein–Barr virus (Henson *et al*., 2009), the Sendai virus (Fujita *et al*., 2006) or the Varicella-zoster virus (Sun *et al*., 2020).

Our work demonstrates that poxvirus infection triggers a five-fold increase in sEVs secretion by immune cells. Similarly, oncolytic poxviruses (including the vaccinia virus) targeting tumor cells were recently shown to increase the secretion of EVs and modify their content (Hirigoyen *et al*., 2024). These findings align with previous reports involving other viral families, such as Enterovirus 71 (Fu *et al*., 2017), rotavirus RRV (Iša *et al*., 2020), hepatitis A virus (Jiang *et al*., 2020), astrovirus (Baez-Navarro *et al*., 2022), and picornavirus (Defourny *et al*., 2024) and supports the notion that increased EVs secretion is a common cellular response to viral infection. While each virus may exploit distinct host pathways, including hijacking the endosomal and MVB pathways, activation of stress and inflammatory responses, and extensive remodelling of the cytoskeleton and membrane trafficking machinery; the convergence on EV-associated pathways, demonstrates a shared strategy of leveraging host cell biology. This evolutionary convergence likely results in altered EV-mediated cell communication from infected cells, which could in turn impact antiviral immunity. While the mechanisms by which poxviruses promote EV secretion are unknown and were not studied here, EV content analysis might provide some hints. We found that sEVs secreted post-infection display reduced surface levels of CD63, a tetraspanin mostly present on multivesicular bodies (MVB) where is sorts cholesterol to nascent exosomes (Palmulli *et al*., 2024). Decrease in CD63 levels, could be related to the capacity of poxvirus to hijack the cellular ESCRT machinery and induce viral maturation in MVBs, as shown for the vaccinia virus (Huttunen *et al*., 2021). We hypothesize that the exploitation of the host machinery responsible for exosome biogenesis by poxvirus diverts the function of the ESCRT complex, thereby decreasing the generation of CD63-positive exosomes. In addition, we observed that EVs secreted post-infection are enriched in actin regulators, including host MYO1A (McConnell et Tyska, 2007), ESPIN (Bartles *et al*., 1996; Rajan *et al*., 2023), profilin and the Arp2/3 complex member ARPC4 (Bieling and Rottner, 2023). This signature might relate to the ability of poxviruses to rapidly remodel actin in order to propel themselves within the cytoplasm (Frischknecht *et al*., 1999; Dodding and Way, 2009; Doceul *et al*., 2010). Since actin remodelling has been associated to EV secretion, both from the plasma membrane (Muralidharan-Chari *et al*., 2009) and from MVBs (Sinha *et al*., 2016), we speculate that poxvirus promote EV secretion by manipulating the actin cytoskeleton. While the precise mechanisms through which poxvirus modulate EV secretion remain to be deciphered, they are likely to involve viral proteins. Among possible candidates, three viral proteins are present in sEVs post-infection and could be involved in boosting EV secretion: B5 and A34, which are known to manipulate host actin cytoskeleton (Wolffe *et al*., 1997; Doceul *et al*., 2010), and F13, which associates to late endosomes (Grosenbach *et al*., 1997; Honeychurch *et al*., 2007). Mapping precisely which EV biogenesis pathways are disrupted, and which subpopulations of EVs are affected by poxvirus infection, will deepen our understanding of virus–host interactions to notably assist in the detection and control of future outbreaks (e.g., mpox, caused by monkeypox virus), using EVs as biomarkers, targets or vaccine vectors. It also reveals opportunities to enhance EV secretion for therapeutic applications, as controlled infection of production cells combined with optimized isolation workflows, could be harnessed to enhance EV yields and support scalable biomanufacturing, thus providing an additional innovative strategy to the methods currently explored (Ng *et al*., 2022).

Manipulation of EV biogenesis by poxviruses might also directly affect their capacity to modulate the immune response. We found that MVA infection of immune cells promotes the accumulation of known mediators of inflammation, either coming from the host cell, such as SAMHD1 (Chen *et al*., 2019), Rack1 (Usacheva *et al*., 2001; Yao *et al*., 2014) or ICAM1 (Bui *et al*., 2020) or from the virus, including B8 (Symons and Smith, 1995) and A46 (Stack and Bowie, 2012). Besides, those EVs also contain viral proteins known to be involved in virus uptake or fusion, such as H3, D8 and B5 (Lin *et al*., 2000; Roberts *et al*., 2009; Pokorny *et al*., 2024) and could therefore have altered internalization route and downstream signaling. More relevant for EV engineering and therapeutic purposes, we identified the transfer of virus-encoded payloads to EVs secreted post-infection: membrane-bound CD40L, the OVA epitope SIINFEKL presented by MHC class I as well as a fraction of the soluble cytokine IL-12 that was co-isolated with sEVs. This offers additional opportunities to amplify the therapeutic efficacy of the vector, for instance by enhancing the enrichment of the payloads in EVs.

Remarkably, we found that EVs secreted by dendritic cells infected with an armed virus and isolated by UC or SEC significantly inhibited tumor growth to levels comparable with an armed virus. This suggest that the efficacy of viral therapies could be partially explained by the release of immunocompetent EVs stimulated by viral infection. Interestingly, the soluble secretome from infected cells similarly decreases tumor growth. While this fraction is deprived of CD40L and contains small amounts of the OVA epitope SIINFEKL presented by MHC class I, it is highly enriched in IL-12 and could contain additional factors, including OVA epitopes, present in protein aggregates/exomeres or in soluble form, able to trigger an anti-tumoral immune response after being processed by APC. Although EVs (UC- and SEC-isolated) and soluble factors similarly reduce tumor growth *in vivo*, it is probable that this is achieved via different mechanisms. Indeed, while EVs isolated by UC induce a strong and direct MHCI-dependent activation and proliferation of OVA-specific T cells *in vitro*, SEC-isolated EVs and soluble fractions only have a moderate effect. Since we detected few but significant amounts of SIINFEKL-specific CD8^+^ T cells in the circulation of mice, it is possible that SEC-isolated EVs and soluble fractions activate T-cells indirectly, via APC, a mechanism that could be amplified by elevated IL-12 levels. Of note, the levels of circulating SIINFEKL-specific CD8^+^ T cells are low compared to the ones found after viral infection, but our analysis was limited to T cells circulating in the peripheral blood, not reflecting what may be occurring in other compartments such as in the tumor microenvironment where higher amount of tumor-infiltrating lymphocytes could be present (Yin *et al*., 2016). Alternatively, SEC-isolated EVs and soluble fractions could promote tumor cell eradication through CD8^+^ T-cell-independent mechanisms, which could include cytotoxic NK cells, known to be activated by IL-12 (Ohs *et al*., 2016; Rademacher *et al*., 2021). The difference between the abilities of the UC- and SEC-isolated EVs to activate CD8^+^ T cells could be explained by the fact that UC co-isolates some co-factors (soluble or aggregated) which are removed by SEC. It is therefore possible that direct T cell activation requires the cumulative effect of both soluble and EV-bound SEC fractions, as shown for instance in the case of the platelet secretome, where both fractions are required to promote angiogenesis and wound closure (Gomes *et al*., 2022). Overall, our work highlights the necessity to precisely characterise the function of each fraction of the secretome, including EVs subpopulations. A better understanding of this complex signaling could open the doors to improved viral vectors further boosting therapeutic cargo loading (Zheng *et al*., 2023; Chen *et al*., 2024).

Altogether our findings demonstrate the anti-tumoral function of sEVs released by immune cells infected with poxviruses. It strengthens the clinical potential of EVs as therapeutics and particularly as vaccine delivery platforms and underscore the importance of considering the entire secretome, as a functionally integrated system capable of driving potent anti-tumor responses.

---

**Table 1: Mass spectrometry on EVs isolated by ultracentrifugation.** Proteins differentially expressed between mock and MVA infected PBMCs are shown (pvalue<0,01). Positive log fold change (logFc): increased expression post-infection. Negative log fold change (logFc): decreased expression post-infection. Data presented on Figure 2 and Supplementary Figure 3.

**Table 2: Mass spectrometry on EVs isolated by ultracentrifugation following 200nm filtration.** Proteins differentially expressed between mock and MVA infected PBMCs are shown (pvalue<0,01). Positive log fold change (logFc): increased expression post-infection. Negative log fold change (logFc): decreased expression post-infection. Data presented on Figure 2 and Supplementary Figure 3.

**Table 3: Mass spectrometry on EVs isolated by ultracentrifugation following 100nm filtration.** Proteins differentially expressed between mock and MVA infected PBMCs are shown (pvalue<0,01). Positive log fold change (logFc): increased expression post-infection. Negative log fold change (logFc): decreased expression post-infection. Data presented on Figure 2 and Supplementary Figure 3.

## Methods

### 1.1. Cell culture

#### Human Primary Cells

Leucocyte concentrates (buffy coats) from healthy donors were provided by the Etablissement Français du Sang (EFS), Strasbourg, France. Human peripheral blood mononuclear cells (PBMCs) were isolated by standard density gradient centrifugation using Ficoll–Paque PLUS (GE Healthcare, Sweden) in 50 mL Blood Separation Tubes (Dacos, Denmark). PBMCs were cultured in RPMI-1640 medium (Sigma) supplemented with 10% heat-inactivated fetal bovine serum (FBS) (#35-070-CV, Corning), 2 mM L-glutamine (Sigma) and 40 µg/mL gentamycin (Sigma) at 37°C and 5% CO_2_. Heat-inactivated FBS was heated for 30 min in a 56°C water bath to inactivate complement.

#### Cell lines

DC2.4 (#SCC142, Merck) are immortalized murine dendritic cells, originally created by transducing bone marrow isolates of C57BL/6 mice with retrovirus vectors expressing murine granulocyte-macrophage CSF (GMCSF) and the myc and raf oncogenes. DC2.4 exhibits characteristic features of dendritic cells, including cell morphology and the expression of dendritic cell-specific markers and the ability to phagocytose and present exogenous antigens on both MHC class I and class II molecules (Shen *et al*., 1997). DC2.4 cells were cultured in RPMI-1640 (Sigma) supplemented with 10% heat-inactivated FCS (#35-070-CV, Corning), 2 mM L-glutamine, 40 µg/mL gentamycin, 1x MEM non-essential amino acids (Gibco), and 1x HEPES Buffer (Gibco) at 37°C and 5% CO_2_.

E.G7-OVA cells (ATCC, CRL-2113^™^) are murine T lymphoblast cell line derived from the C57BL/6 (H-2 b) mouse lymphoma cell line EL4. E.G7-OVA cells were cultured in RPMI-1640 medium (Sigma), supplemented with 10% heat-inactivated FCS (#35-070-CV, Corning), with 2 mM L-glutamine adjusted to contain 1.5 g/L sodium bicarbonate, 4.5 g/L glucose, 10 mM HEPES and 1.0 mM sodium pyruvate and supplemented with 0.4 mg/mL G418, (all provided by Sigma) at 37°C and 5% CO_2_.

U-2OS cells (ATCC, HTB-96^™^) are derived in 1964 from a moderately differentiated osteosarcoma of the tibia of a 15-year-old, White, female patient, they were cultured in DMEM (Gibco), supplemented with 10% FCS (#35-070-CV, Corning), 2 mM L-glutamine (Sigma) and 40 µg/mL gentamycin (Sigma) at 37°C and 5% CO_2_.

BHK-21 cells (ATCC, CCL-10^™^) are fibroblasts isolated from the kidney of an uninfected golden hamster, they were cultured in DMEM High Glucose (Gibco), supplemented with 10% FCS (#35-070-CV, Corning) at 37°C and 5% CO_2_.

BT cells (CRL-1390^™^) are bovine *Bos Taurus* turbinate cells, they were cultivated in DMEM supplemented with 10% fetal horse serum (Hyclone, Logan, USA), 2 mM L-glutamine (Sigma) and 40 mg L^-1^ gentamicin (Sigma). Regular mycoplasma testing was carried out for all cell lines (Clean Cells, Vendée, France or Taconic Biosciences).

### 1.2. Viral vectors

All viruses were generated by homologous recombination and amplified in chicken embryo fibroblast (CEF) for VV_COP_ and MVA or in BT cells for PCPV. Cells were transfected with the shuttle plasmids encoding the expression cassettes (described below) and previously infected with parental virus. Recombinant viruses were isolated by picking non-fluorescent viral plaques. The identity and integrity of expression cassettes were checked by PCR analysis and DNA sequencing of each locus. All viruses were purified by tangential flow filtration and titrated by plaque assay.

The following vectors were previously described: VV_COP_-eGFP (Foloppe *et al*., 2019). MVA-eGFP and ^empty^MVA (Fournillier *et al*., 2007; Erbs *et al*., 2008), PCPV-eGFP (Ramos *et al*., 2022). OVA MVA, OVA-IL12 MVA, OVA-CD40L MVA and OVA-CD40L-IL12 (corresponding to the ^armed^MVA), shown in Fig.S4, encode their respective payloads: a polyepitope cassette comprising two MHC I–restricted epitopes (SIINFEKL, OVA_257-264_/H-2Kᵇ; WMHHNMDLI/H-2Dᵇ) and two MHC II–restricted epitopes (ISQAVHAAHAEINEAGR, OVA_323-339_/I-Aᵇ; NAGFNSNRANSSRSS/I-Aᵇ) fused to eGFP under the pH5R promoter in deletion III, murine CD40L under the pH5R promoter in deletion II and murine IL-12 under the early/late 11K7,5 promoter in deletion II.

### 1.3. Cell culture conditions for EVs production

PBMCs were cultured in complete medium for testing the various isolation methods described with VV_COP_-eGFP. When using MVA-eGFP with PBMCs for mass spectrometry analysis of EVs content and all experiments involving DC2.4, cells were cultured in EV-depleted medium that was prepared using proper 2x complete medium, ultracentrifuged at 100,000 g for 18h (Beckman Coulter Optima XL-80K Ultracentrifuge with a SW28 rotor) to remove EVs from FBS. The resulting supernatant was adjusted to 1× with the appropriate serum-free medium for each cell type.

### 1.4. Cell infection

PBMCs were infected with virus of interest at MOI=1 and incubated for 18h, and DC2.4 were infected with virus of interest at MOI=1 and incubated for 18h, before harvesting the supernatant containing EVs.

### 1.5. EVs isolation and quantification

After infection and incubation, supernatant was pre-cleared by centrifugation at 300 g for 10min and 2,000 g for 30min, at room temperature.

#### Ultracentrifugation (UC)

Pre-cleared supernatant was centrifuged at 10,000 g for 45min at room temperature before a final ultracentrifugation (Beckman Coulter Optima XL-80K Ultracentrifuge in 38.5 mL Open-Top Thinwall Polypropylene tubes with a SW28 rotor, Beckman Coulter) at 100,000 g for 2h, at 4°C. The supernatant was discarded and resulting EVs pellet was resuspended in PBS and stored at 4°C.

#### Immunocapture

Pre-cleared supernatant was centrifuged at 10,000 g for 45min at room temperature and EVs were isolated using the EV Isolation Kit Pan (#130-110-912, Miltenyi Biotec), according to the manufacturer’s instructions. Two fractions were obtained: the FT, corresponding to the flowthrough fraction containing non-bound components to MicroBeads, and the CAPT. fraction, corresponding to eluted EVs (and other co-purified particles) bound to MicroBeads and retained prior elution.

#### Iodixanol density gradient

Pre-cleared supernatant was concentrated using a Centricon Plus-70 10kDa (#UFC701008, Merck Millipore) according to the manufacturer’s instructions. An iodixanol gradient (Optiprep^TM^ 60% iodixanol (w/v), Serumwerk Bernburg, Germany) was used as described by Van Deun *et al*., 2014. Briefly, the gradient was formed by gently layering 4 mL of 40%, 4 mL of 20%, 4 mL of 10% and 3.5 mL of 5% iodixanol solutions on top of each other in a 17 mL polypropylene tube (Beckman Coulter). 1 mL of concentrated sample was overlaid on the top of the gradient which was then centrifuged for 18 hours at 100,000 g and 4°C (Beckman Coulter XL-70 centrifuge with a SW 32.1 Ti rotor). To prevent disturbance of gradient, deceleration was performed without applying brakes. 16 fractions of 1 mL and 1 fraction of 0.5 mL were collected from the top to the bottom of the gradient, diluted in PBS and centrifuged for 3 hours at 100,000 g and 4°C. After discarding the supernatant, the resulting pellets were resuspended in 100 µL PBS and stored overnight at 4°C before being analysed by NTA and titration assays.

#### Iodixanol velocity gradient

Pre-cleared supernatant was concentrated using a Centricon Plus-70 10kDa (#UFC701008, Merck Millipore) according to the manufacturer’s instructions. An iodixanol gradient (Optiprep^TM^ 60% iodixanol (w/v), Serumwerk Bernburg, Germany) was used as described by Martin-Jaular *et al*., 2021. Briefly, each solution was prepared by diluting Optiprep^TM^ in PBS to the indicated iodixanol concentration, and the gradient was formed by gently layering 3 mL of 18%, 3 mL of 14%, 3 mL of 10% and 2.5 mL of 6% iodixanol solutions on top of each other in a 13 mL polyallomer tube (Beckman Coulter). 1 mL of concentrated sample was overlaid on the top of the gradient which was then centrifuged for 1 hour at 200,000 g and 4°C (using a Beckman Coulter XL-70 centrifuge with a SW 32.1 Ti rotor). To prevent disturbance of gradient, deceleration was performed without applying brakes. 12 fractions of 1 mL and 1 fraction of 0.5 mL were collected from the top to the bottom of the gradient, diluted in PBS and centrifuged for 45min at 100,000 g and 4°C. After discarding the supernatant, the resulting pellets were resuspended in 100 µL PBS and stored overnight at 4°C before being analysed by NTA and titration assays.

#### Filtration

Pre-cleared supernatant was concentrated using a Centricon Plus-70 10kDa (#UFC701008, Merck Millipore) at room temperature, according to the manufacturer’s instructions. Concentrated samples remained unfiltered or were passed through filters with 200nm (0,2µm, SFCA, 25mm, #723-2520, Nalgene) or 100nm (Millex 0,1 µm, PVDF, 33 mm**, #**SLVV033RS, Merck Millipore) pores and EVs were either recovered by UC or SEC afterward.

#### Size Exclusion Chromatography (SEC)

When tested as a standalone isolation method, the first generation of qEV2 70nm Legacy columns (Izon Science Ltd, New Zealand) were used according to the manufacturer’s instructions. In brief, pre-cleared supernatant was concentrated using a Centricon Plus-70 10kDa (#UFC701008, Merck Millipore). After rinsing the column with PBS, 2 mL of concentrated sample were loaded before adding 12 mL of PBS; this void volume was discarded. Then, 2 mL of PBS were successively added to harvest ten fractions of 2 mL. Each fraction was stored overnight at 4°C before being analysed by NTA and titration assays.

When used to recover EVs after 100nm filtration and enhance their separation from soluble proteins, the qEV2 35nm Gen 2 columns (Izon Science Ltd, New Zealand) were used according to the manufacturer’s instructions. After rinsing the column with PBS, 2 mL of concentrated sample were loaded before adding 13.9 mL of PBS; this void volume was discarded. Then, 2 mL of PBS were successively added to harvest ten fractions of 2 mL. Fractions 1 to 4 were pooled to form the EVs fraction and fractions 5 to 10 were pooled to form the proteins-enriched fraction, before being concentrated using Amicon Ultra-15 Centrifugal Filter Unit 10 kDa (#UFC901024, Merck Millipore). The volume of both samples was equalized with PBS, analysed by NTA and stored at 4°C.

Unless otherwise stated, EVs for functional assays and in vivo experiments were quantified by NTA and either used immediately after isolation or stored overnight at 4 °C; stored samples were analysed or injected the following day. For proteomic analyses by mass spectrometry EVs were quantified by NTA, aliquoted, and stored at −80 °C until analysis.

#### Nanoparticles Tracking Analysis (NTA)

According to the manufacturer’s procedure, EVs (or particles) were diluted in PBS and the number of EVs (or particles) was determined using the ZetaView analyser (PMX-120) from Particle Metrix GmbH (Meerbuch, Germany). The sensitivity of the camera was set to 80 for all measurements. The data were analysed using the ZetaView analysis software version 8.05.16 SP7. The quantities of EVs (or particles) are expressed as the number of EVs (or particles)/secretory cells, counted when seeding the culture flasks just before infection.

### 1.6. Infectivity titration

Samples were titrated by plaque assays, performed on virus-specific indicator cells: U-2 OS (samples from VV_COP_–infected PBMCs), BHK-21 (samples from MVA-infected PBMCs or DC2.4), and BT cells (samples from PCPV-infected PBMCs). Confluent monolayers in 6-well plates were inoculated with 10-fold serial dilutions of samples (duplicate wells), adsorbed for 30min to 2h at RT or 37 °C (depending on the cell type) prior addition of medium containing agarose or carboxymethyl cellulose. Plates were incubated until plaques were countable (72h for VV_COP_; 30h for MVA; 48h for PCPV). Plaques were visualized by neutral red (PCPV) or crystal violet staining (VV_COP_); or were revealed by immunostaining with an anti-vaccinia virus antibody (#B65101R, Meridian Bioscience) followed by an HRP-conjugated secondary (#P0448, Dako) and development with a chromogenic peroxidase substrate (MVA). PFU/mL were calculated from dilutions yielding 20–100 plaques.

### 1.7. Mass spectrometry

#### EVs samples preparation

As described above, EVs were isolated by UC from PBMCs (cultured in EVs-free medium) that were either uninfected (mock) or infected with Modified Vaccinia Ankara (MVA) encoding eGFP at MOI=1. Four independent replicates were performed per condition (6 conditions; n = 24 total samples). For proteomics, 1×10⁹ particles per sample were collected and stored at −80 °C until mass-spectrometry analysis.

#### Electrophoresis and in gel trypsin digestion

Samples were lysed using 135 µl of Laemmli buffer. The whole material was loaded on an in-house poured 4% acrylamide stacking gel. Gel was stained with Coomassie Blue and the stacking bands were manually excised. Proteins were then reduced using 10 mM dithiothreitol and alkylated using 55 mM iodoacetamide before in-gel digestion overnight at 37°C with 0.2 µg of trypsin/LysC (Promega, Madison, USA). Peptides were extracted for 1 h with 90 μl of 80% acetonitrile, 0.1% formic acid. Peptide mixtures were then dried and resuspended in 50 µl water acidified with 0.1% formic acid.

#### Liquid Chromatography-Tandem Mass Spectrometry (LC-MS/MS) Analyses

LC-MS/MS analyses of peptide extracts were performed on a NanoElute LC-system coupled to a TimsTOF Pro mass spectrometer equipped with a CaptiveSpray ion source (Bruker Daltonics, Billerica, MA, USA). Mobile phase A was constituted of 97.9% water, 2 % acetonitrile and 0.1% formic acid and mobile phase B was constituted of 99.9% acetonitrile and 0.1% formic acid. A sample volume of 1 µl was loaded into an AcclaimPepMap C18 precolumn (0.1 x 20 mm, 5 μm particle size, ThermoFisher Scientific) at 120 bars and 2% B. This step was followed by reverse-phase separation at a flow rate of 300 nl/min using an Aurora 2 C18 separation column (250 mm x 75 μm id, 1.6 μm particle size, IonOpticks) and a gradient from 2% to 25% B in 90 min, from 25% to 35 % B in 10 min, from 35% B to 85% B in 13 min, and maintained at 85% B for 17 min. Precolumn and column were further reconditioned using 10 or 5 volumes of 2% B respectively.

The TimsTOF Pro instrument was operated in Data Dependent Acquisition-Parallel Accumulation Serial Fragmentation (DDA-PASEF) mode with 10 PASEF scans in a mass range from 100 m/z to 1700 m/z leading to 1.17 s cycle time. The ion mobility range scanned from 0.7 to 1.25 V.s/cm² with accumulation and ramp times of 100 ms. Peptides with a minimum intensity of 2500 were selected for fragmentation and were automatically added on a dynamic exclusion list for 0.4 min. Collisional induced dissociation (CID) voltage was tuned according to the mobility of the selected ion using a ramp from 20 eV for 0.6 V.s/cm² to 59 eV for 1.6 V.s/cm².

#### LC-MS/MS data interpretation and validation

Raw files were converted to .mgf peaklists using Data Analysis (version 5.3, Bruker Daltonics) and were submitted to Mascot database searches (version 2.6.2, MatrixScience, London, UK) against a multispecies protein database (total of 108,981 entries) including *Homo sapiens* and Modified Vaccinia Ankara virus sequences as target sequences, plus *Bos Taurus* and *Gallus gallus* as contaminant sequences, all extracted from UniProtKB-SwissProt and UniProtKB-TrEmbl. Benjamini Hochberg strategy was used to validate true identifications (Couté *et al*., 2020). Spectra were searched with a mass tolerance of 10 ppm in MS mode and 0.05 Da in MS/MS mode. One trypsin missed cleavage was allowed. Carbamidomethylation of cysteine residues was set as a fixed modification. Oxidation of methionine residues and acetylation of protein n-termini were set as variable modifications. Identification results were imported into Proline software (http://proline.profiproteomics.fr/; Bouyssié *et al*., 2020) for validation. Peptide Spectrum Matches (PSM) with pretty rank equal to one and a minimum length of 7 amino acids were retained. False Discovery Rate was then optimized to be below 1% at PSM level using Mascot score and below 1% at Protein Level using Fisher score.

#### Label Free Quantification

Peptide abundances were extracted thanks to Proline software version 1.0 (http://proline.profiproteomics.fr/; Bouyssié *et al*., 2020) using a m/z tolerance of 10 ppm. Alignment of the LC-MS runs was performed using Loess smoothing. No Cross assignment was allowed. The best 2+/3+/4+ peptide ion was used to assign an abundance to a peptide. Protein abundance was then computed by summing the abundance of all specific peptides.

#### Statistical Analysis

Protein abundances of human and viral proteins (2050 proteins) were loaded into Prostar software version 1.26.4 (http://www.prostar-proteomics.org/; Wieczorek *et al*., 2017) and associated to the 6 conditions (mock no filtration, mock filtered 200nm, mock filtered 100nm, MVA no filtration, MVA filtered 200nm, MVA filtered 100nm). Proteins with at least 3 values in at least one condition (3/4 replicates) were kept for further statistical analysis. Protein abundances were normalized using 15% quantile centering. Residual missing values were imputed in a conservative way (quantile 2.5%). Pairwised Student t-tests were performed to compare mock vs MVA infected cells according to the three filtration statuses.

P-values calibration was corrected using Benjamini-Hochberg method, and FDR was set to ∼1-4%. More precisely, the comparison mock vs MVA without filtration leads to 0.95% FDR using p-values below 0.00209, the comparison mock vs MVA with 0.2 µm filter leads to 3.93% FDR using p-values below 0.00229, the comparison mock vs MVA with 0.1 µm filter leads to 2.63% FDR using p-values below 0.00182.

The mass spectrometry proteomics data have been deposited to the ProteomeXchange Consortium via the PRIDE partner repository (Perez-Riverol *et al*., 2022).

Proteins with p < 0.01 and log2FC ≥ 0.5 were retained for heat maps and volcano plots, generated with custom R functions and the R package Pheatmap. GO term analysis was performed using https://string-db.org/ and all proteins with -log(pValue) >1,3 were considered.

### 1.8. EV characterisation

#### EV surface marker profiling by electrochemiluminescence (ECL) immunoassay

Assays were performed on MSD 1-Spot High Binding SECTOR Plate (L15XB, Meso Scale Discovery) following the manufacturer’s instructions. Wells were coated with 25 µL/well of samples solution, containing 2,5.10^8^ EVs. Plates were sealed and incubated 2h at room temperature (RT) without shaking. Plates were gently washed with 150 µL PBS/well, then blocked with 150 µL/well of PBS 5% Blocker A (R93BA, Meso Scale Discovery) for 1h at RT with gentle orbital shaking. After a washing with 150 µL PBS/well, 25 µL/well of detection antibody solution (diluted at 1 μg/mL in PBS 5% Blocker A) were added and incubated for 2h at RT with gentle orbital shaking. Plates were then washed 3 times with 150 µL PBS/well, incubated with 25 µL/well SULFOTAG Streptavidin (R32AD, Meso Scale Discovery) for 30min at RT with gentle orbital shaking, then washed three times with 150 µL PBS/well. Finally, 150 µL/well of MSD GOLD Read Buffer B (R60AM, Meso Scale Discovery) were added and plates were read immediately on a MESO QuickPlex SQ 120MM instrument. The following detection antibodies were used: CD63 Antibody, anti-mouse Biotin (clone REA563, Miltenyi Biotec); CD54 (ICAM-1) Antibody, anti-mouse, Biotin (clone YN1/1.7.4, #130-104-213, Miltenyi Biotec); CD86 Antibody, anti-mouse, Biotin (clone PO3.3, #130-101-944, Miltenyi Biotec); CD154 (CD40L) Antibody, anti-mouse, Biotin (clone MR1, #130-101-900, Miltenyi Biotec); H-2Kb/SIINFEKL Antibody, anti-mouse Biotin (clone 25-D1.16, Miltenyi Biotec).

#### IL-12p70 quantification by electrochemiluminescence (ECL) immunoassay

Il-12p70 was quantified using the V-PLEX Proinflammatory Panel 1 mouse kit (K152QVD, Meso Scale Discovery), following the manufacturer’s assay protocol. Briefly, MULTI-SPOT 96-well plate pre-coated with capture antibodies was washed 3 times with 150 µL/well of wash buffer (R61AA, Meso Scale Discovery). Samples, calibrators, or controls (50 µL/well) were added, plates were sealed and incubated 2 h at room temperature (RT) with gentle orbital shaking. Plates were then washed 3 times with 150 µL/well of wash buffer, 25 µL/well of the SULFO-TAG–conjugated IL-12p70 detection antibody solution (D22QV, Meso Scale Discovery) was added, and plates were incubated 2h at RT with gentle orbital shaking. After the final three washes with 150 µL/well of wash buffer, 150 µL/well of MSD GOLD Read Buffer B (R60AM, Meso Scale Discovery) were added and plates were read immediately on an a MESO QuickPlex SQ 120MM instrument. Raw signals (ECL units) were converted to concentrations using the kit’s calibration curve and analysed with the MSD software.

### 1.9. OT-I T cells isolation and activation

All animals were housed and handled at INSERM (mouse facility agreement number: #C67-482-33) according to the guidelines of INSERM and the ethical committee of Alsace (CREMEAS), following French and European Union animal welfare guidelines (Directive 2010/63/EU on the protection of animals used for scientific purposes). All procedures were performed in accordance with the ARRIVE guidelines, French and European Union animal welfare guidelines and supervised by local ethics committee. Animals were housed in pathogen-free conditions with food and water ad libitum and appropriate enrichment (sterile pulp paper and coarsely litter).

Single-cell suspensions of splenocytes were obtained by mechanical disruption of spleen from 8–12-week-old female wild-type C57BL/6-Tg (TcraTcrb)1100Mjb/Crl OT-I TCR transgenic mice (642OT1; Charles River) through a 30µm nylon cell strainer (MiltenyiBiotec #1130-098-458). Red blood cells were lysed in ACK Buffer [150mM NH_4_Cl, 10mM KHCO_3_, 0.1mM Na_2_EDTA] for 3 min at room temperature. Naïve CD8^+^ T cells were purified with a mouse Naïve CD8a^+^ T cell Isolation Kit (MiltenyiBiotec #130-096-543). Purified naïve CD8^+^ T cells purity was assessed by flow cytometry on a Attune NxT (Invitrogen) flow cytometer using FITC anti-CD4 (1:100, clone RM4-5, BioLegend #100510), BV605 anti-CD8 (1:200, clone 53-6.7, BioLegend #100743), PE anti-CD44 (1:200, clone IM7, BioLegend #103024), PE-Cy7 anti-TCR Va2 (1:100, clone B20.1, BioLegend #103024), and APC anti-TCR Vb5.1, 5.2 (1:100, clone MR9-4, BioLegend #139506) staining and ranged from 95.7 to 99.8 (median 99.3%).

Purified naïve CD8^+^ T cells were then stained with 0.5µM CellTrace Violet dye (Invitrogen #C34571) at 1×10^7^ cells/ml in PBS for 15 min at 37°C in the dark before quenching with 1 vol ice-cold FCS for 5 min on ice. Labelled cells were washed, and 2×10^5^ cells were treated with 5×10^9^ EVs (i.e. 2,5×10^4^ EVs/cell) at 37°C/5% CO_2_ for 72h in flat-bottomed 96 well plates in RPMI 1640 (Gibco #72400054) containing 2mM Glutamine, 25mM HEPES and completed with 10 % v/v of FBS (Gibco), 100 U/mL penicillin, 100 μg/mL streptomycin (PanBiotech #P06-07100) and 50 μM b-mercaptoethanol (Gibco #11508916). For positive control, 2×10^5^ cells were cultured as above in wells previously coated with 5µg/mL of anti-CD3e (clone 145-2C11, BioLegend #100301) and 5µg/mL of anti-CD28 (clone 37.51, BioLegend #102101) in PBS overnight at 4°C. After incubation, the cells were washed in PBS and non-viable cells were stained with Fixable Viability Dye-eFluor780 (1:1,000, Invitrogen #L34993) for 15 minutes at room temperature in the dark. Then, cells were stained with PE anti-CD44 (1:200, clone IM7, BioLegend #103024) for 15 min at 4°C. Samples were acquired with Attune NxT (Invitrogen) flow cytometer and data were analysed using Kaluza Analysis Software 2.4 (Beckman Coulter).

### 1.10. Mice and E.G7-OVA tumor model

All animals were housed and handled at the “Institut Clinique de la Souris” (mouse facility agreement number: #C67-218-40). All procedures experiments were approved by the local ethical committee (C2EA - 17 Comité d’éthique Com’Eth) for Animal Care and Use and research minister (APAFIS #40516 and #36730).

Female C57BL/6 and BALB/c mice 7 to 8 weeks of age were used for the experiments. The mice were purchased from Charles River and maintained in a temperature- and humidity-controlled animal facility (21+/- 2°C; 15-70% humidity), with a 12-h light-dark cycle and free access to water and a standard rodent chow (D04, SAFE, Villemoisson-sur-Orge, France).

E.G7-OVA cells were subcutaneously injected at 3×10^5^ cells per mouse in one abdominal flank. Therapeutic vaccines (1×10^7^ PFU/mice) or EVs (9×10^8^ to 4×10^9^ EVs/mice) were injected intravenously 7 and 14 days after tumor inoculation. The size of superficial tumor was assessed three times per week by using a caliper and the body weight (BW) was monitored at the same time. Tumor volume was calculated with the spheroid formula V = (LxW^2)/2 after calculation of Width and Length diameter in the right angles. A maximal tumor volume of 2000 mm^3^ and a maximum of 10% of BW loss between two measures have been considered as an ethical endpoint. Additional endpoint like ulceration, necrosis, and distension of covering tissues are recorded and would lead to terminate animals humanely when the degree of suffering cannot be justified by the scientific objective.

### 1.11. Ex vivo studies / Pentamer SIINFEKL staining

To detect and quantify CD8^+^ OVA-specific T cells, a MHC-pentameric assay was used. Lymphocytes were isolated from heparinized blood taken at day D21 after tumor inoculation and incubated with 10 µL/test of R-PE Pro MHC Pentamer H-2K^b^ SIINFEKL (#F093-2C-E, ProImmune) at +4°C for 30 min. Cells were resuspended and stained in wash buffer containing CD8a APC (1:50, clone 53-67, BD Pharmingen #553035) and CD19 V450 (1:50, clone 1D3, Tonbo biosciences #75-0193-U100) antibodies and incubated for 30 min at +4°C before flow cytometry analysis. The Pentamer-positive cells were viewed by gating first on CD19 negative lymphoid cells and then analysing on a two-color plot showing CD8 vs Pentamer. Samples were acquired with Aurora (Cytek) flow cytometer and data were analysed using SpectroFlo^®^ (Cytek) software.

### 1.12. Cryo-electron microscopy

Sample vitrification was performed using grids with a holey carbon film (Quantifoil R2/2) to which an additional continuous carbon support was added. Grids were cleaned for 1 minute 30 in a Fischione 1070 plasma cleaner, in an 80% Argon/20% Oxygen gas mix. 3 µL of sample were applied on the grids, and excess of liquid was removed after 30s of incubation by blotting for 5s before vitrification by fast plunging in liquid ethane, using a Vitrobot Mark IV (Thermo Fisher Scientific) setup at 15°C and 95% humidity. Grids were observed on the Glacios TEM (Thermo Fisher Scientific) operated at 200 kV, and images were recorded on K2 camera (Gatan) with a pixel size of 1.9A/pixel and a total dose of about 20e/A2 per image.

### 1.13. Statistical analyses

Normality was determined using a Shapiro-Wilk test. Unpaired t test was used for paired comparison data following a normal distribution and Mann-Whitney test otherwise. Multiple comparison was analysed using a one-way anova or a two-way anova test for data following a normal distribution or a Kruskal-Wallis test otherwise.

## Authors Contributions

LW designed, performed and analysed most of the experiments. NS designed and provided the viruses. MCC and KR helped with EV isolation and SIINFEKL pentamer staining. VM initiated and performed the *in vitro* T cell experiments with help from LB and AL. MR and CCa were responsible for the mass spectrometry, with help from JD and VH for analysis. JGG, KR and VH supervised the project, designed and analysed the experiments. EQ enabled the implementation and development of the project throughout. LW, JGG, KR and VH wrote the manuscript.

## Acknowledgments and fundings

We thank members of the Tumor Biomechanics Lab and Transgene for helpful discussions. We thank Kuang-Jing Huang for helping with electron microscopy pictures processing. We warmly thank Xavier Huet from Meso Scale Discovery (MSD) for helping with ECL immunoassays data processing. We thank Carine Reymann and Yasmin Schlesinger for the generation of recombinant MVA. We thank members of the CRBS animal facility for animal care and the members of the “Institut Clinique de la Souris” (Illkirch-Graffenstaden, France) who handle the E.G7-OVA *in vivo* experiments. We thank Alexandre Durand and Corinne Crucifix from the IGBMC (“Institut de Génétique et de Biologie Moléculaire et Cellulaire”, Department of Integrative Structural Biology, Illkirch-Graffenstaden, France) for performing cryo-electron microscopy and image acquisition. Research was co-funded by Transgene S.A. and a CIFRE PhD fellowship to LW managed by the Association Nationale de la Recherche et de la Technologie, ANRT (grant n°2021/0562). VM was supported by a fellowship from the French Ministry of Science (MESRI) and a fourth-year thesis fellowship from the Fondation ARC pour la recherche sur le cancer (n° ARCDOC42023010006005). LB was funded by PhD fellowship from Fondation pour la Recherche Médicale (n°ECO202206015567). Proteomics analyses were supported by the French National Research Agency ProFI fundings (Proteomics French Infrastructure; ANR-10-INBS-08; ANR-24-INBS-0015). This work was also supported by grants from La Ligue contre le Cancer and by institutional funds from University of Strasbourg and INSERM.

---

**Supplementary Figure 1:**
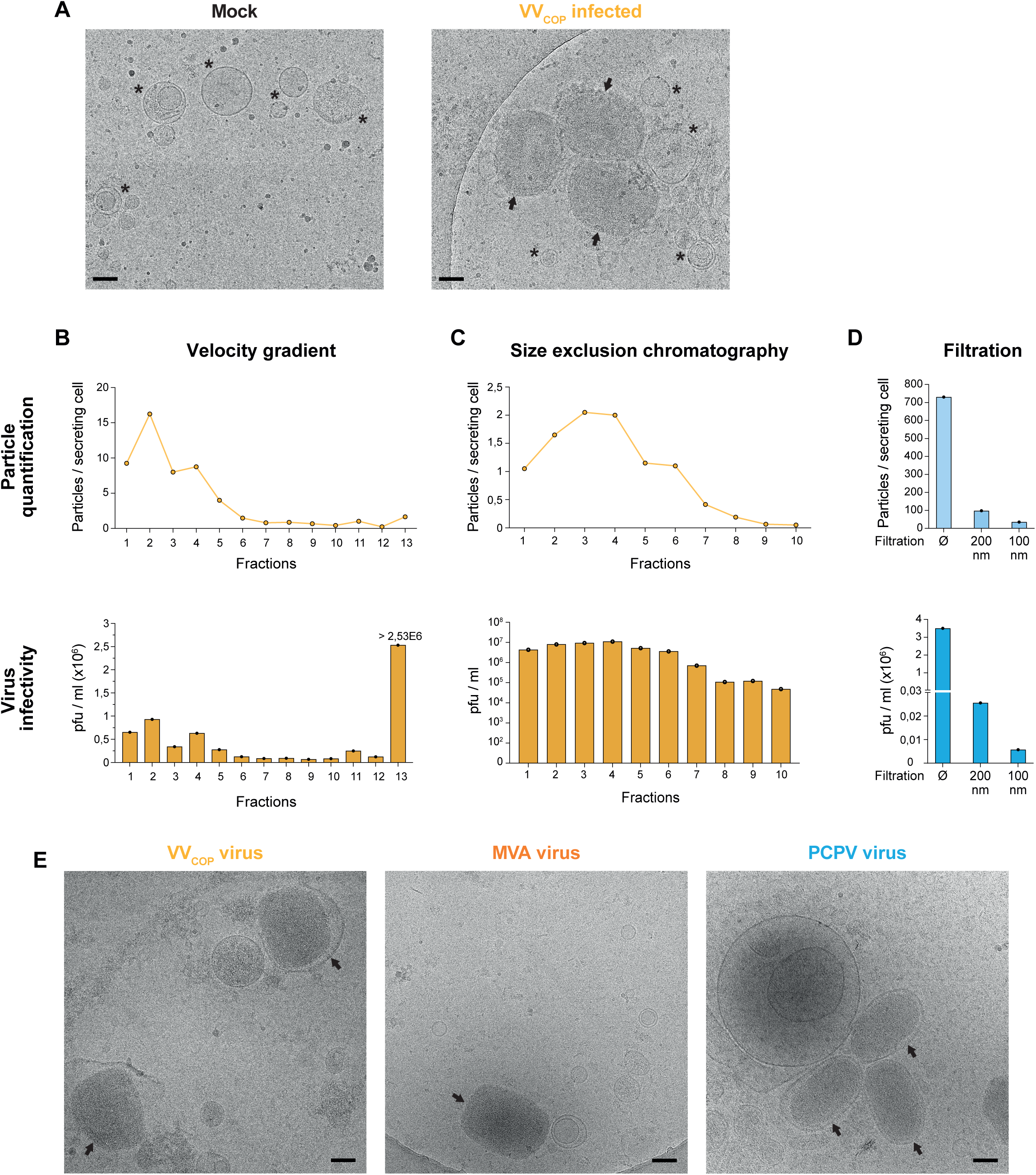
Ultracentrifugation, velocity gradient and size exclusion chromatography do not separate EVs from infectious poxvirus particles. A) Cryo-EM images of particles retrieved in 100k g pellet after ultracentrifugation of culture medium from non-infected (left) or VV_COP_-infected human peripheral blood mononuclear cells (PBMC), showing the presence of EVs (black stars) and electron dense viral particles (black arrows). Scale bar: 100nm. B-C) EVs were isolated either by velocity gradient (B) or by size exclusion chromatography (C) from the supernatant of non-infected PBMCs (mock) or infected by VV_COP_ at a multiplicity of infection of 1 (MOI=1). In each fraction, particles were quantified by NTA (top graphs) and virus infectivity was assessed by titration of plaque forming units (bottom graphs). N=1 experiment each. D) 100nm filtration coupled to ultracentrifugation is not able to fully remove PCPV infectious particles. E) Cryo-EM images of stocks from VV_COP_, MVA and PCPV viruses showing the shape of viral particles (black arrows). Scale bar: 100nm.

**Supplementary Figure 2:**
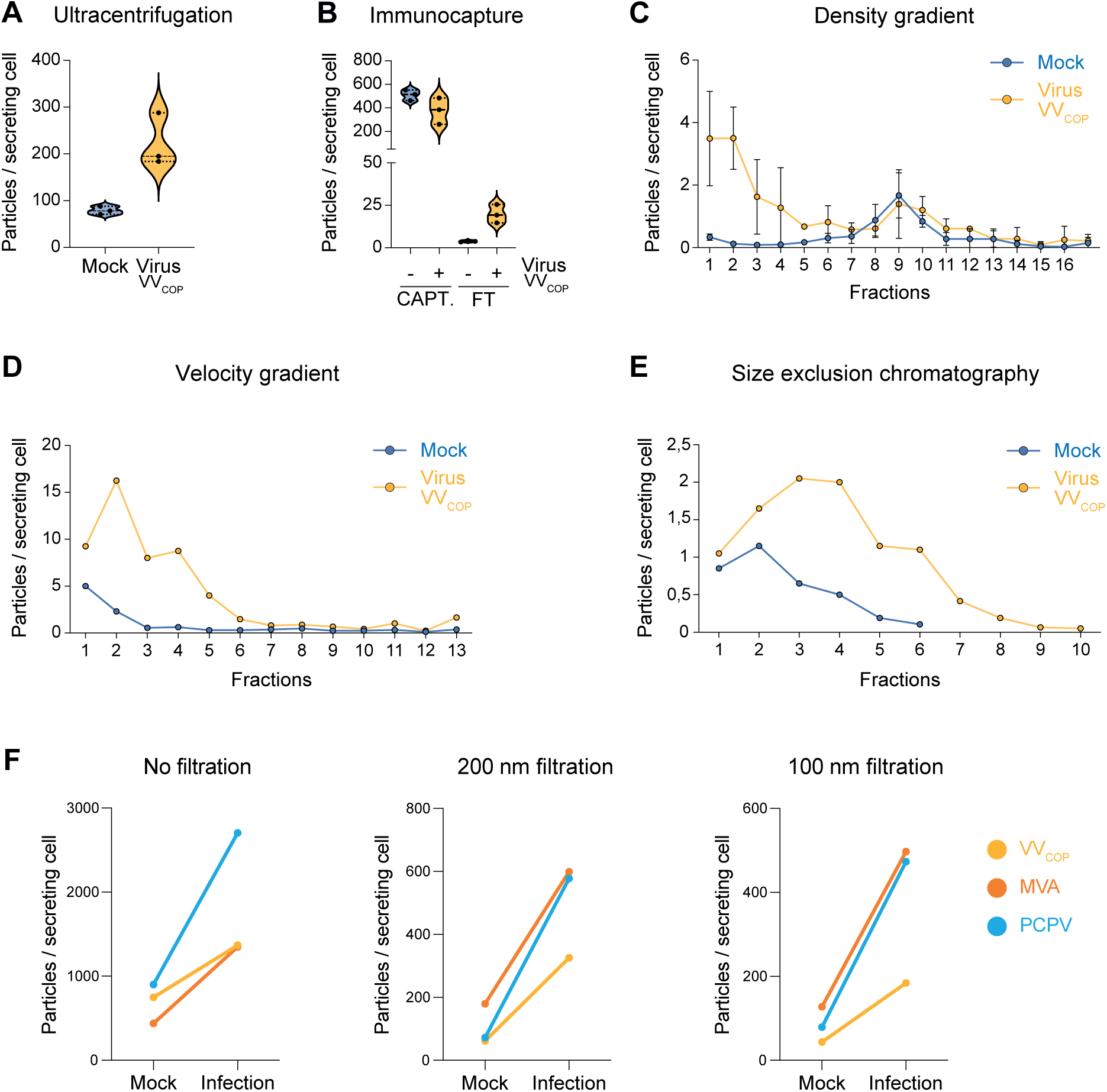
Poxvirus infection increases sEVs secretion. A-E) NTA quantification of EVs and particles isolated by ultracentrifugation (A), immunocapture (B), density gradient (C), velocity gradient (D) or size exclusion chromatography (E) from non-infected PBMCs (mock, blue) or infected with VV_COP_ virus (yellow). F) NTA quantification of EVs isolated by ultracentrifugation and filtered at 200nm or 100nm from PBMCs infected with either VV_COP_ (yellow), MVA (orange) or PCPV (light blue) virus.

**Supplementary Figure 3:**
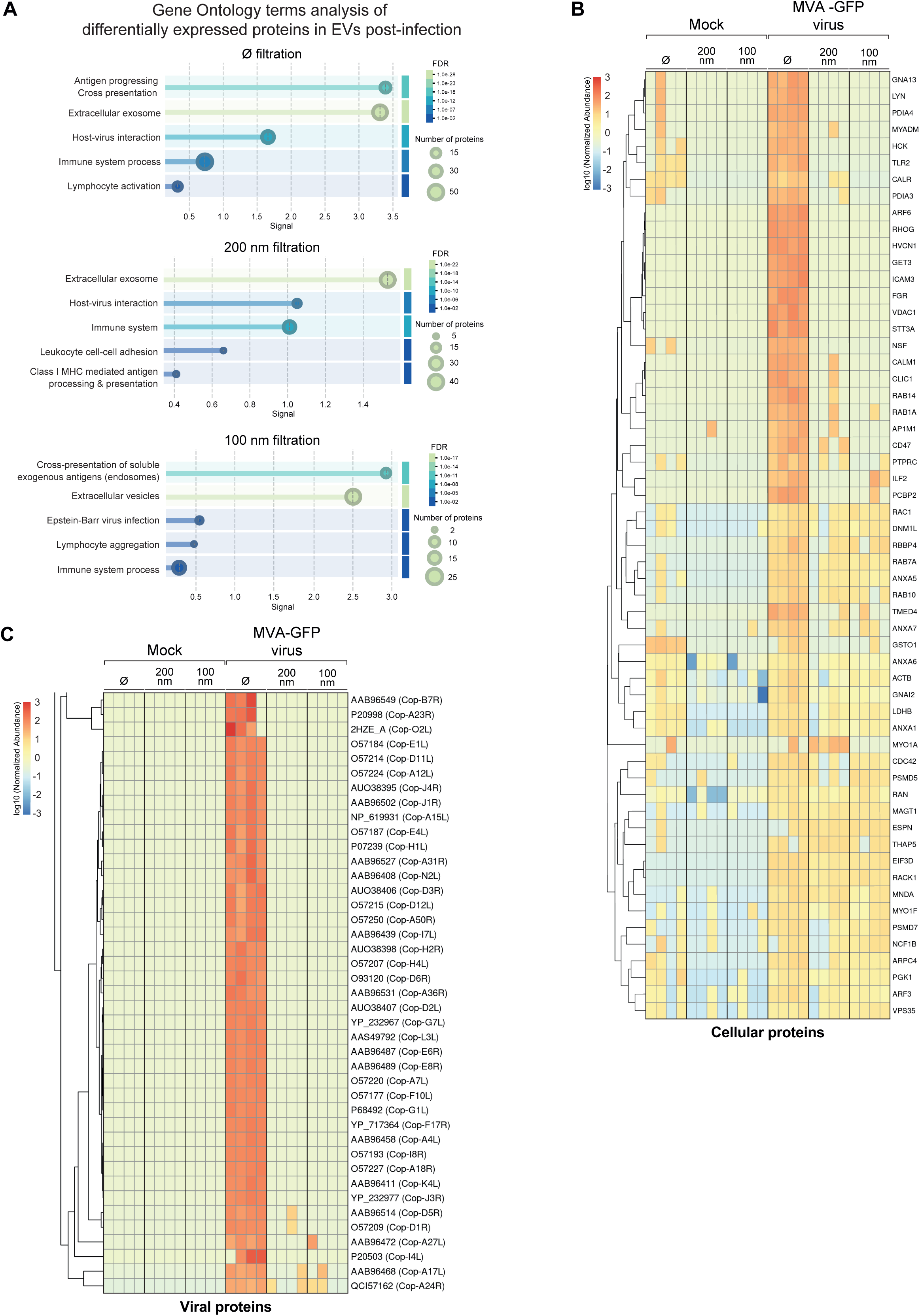
Poxvirus infection alters the content of secreted EVs. Mass spectrometry analysis of the protein content of EVs secreted by mouse dendritic cells (DC2.4) infected by MVA poxvirus and isolated by ultracentrifugation preceded or not by 100nm or 200nm filtration. Mass spectrometry was performed in quadruplicate. A) Gene ontology analysis of the EVs proteins differentially expressed upon poxvirus infection. B-C) Heat maps showing differentially expressed cellular (B) and viral (C) proteins identified in ultracentrifugation pellet with or without filtration (100nm or 200nm). In C), label corresponds to MVA Uniprot ID (VVcop ortholog). The heat map of viral proteins complements the heat map in Figure 2E and applies the same color code, log10 transformation, and normalization of abundance values.

**Supplementary Figure 4:**
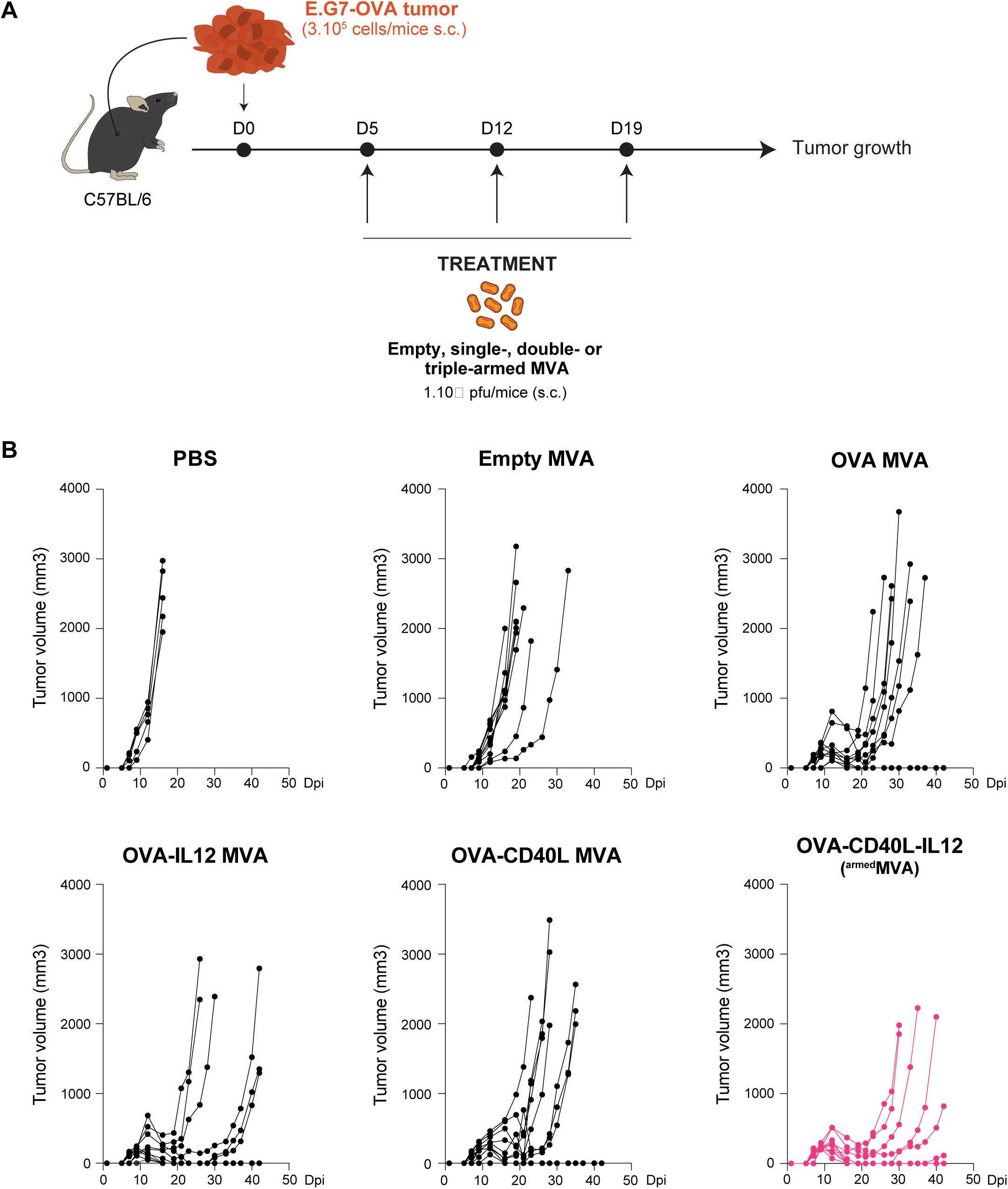
Multi-armed therapeutic poxvirus impairs tumor growth. A) Protocol used to determine the anti-tumoral potential of MVA virus encoding different therapeutic payloads. Viruses were injected intravenously 5-, 12- and 19-days post E.G7-OVA tumor cell injection and tumor growth was monitored for 42 days. B) tumor growth curves of individual mice treated with PBS, with empty virus or with virus encoding the OVA antigens alone or in combination with IL-12, CD40L or both. Arrows indicate the injection days. One curve represents one mouse, n= 5 to 10 mice per group, in N=1 experiment.

**Supplementary Figure 5:**
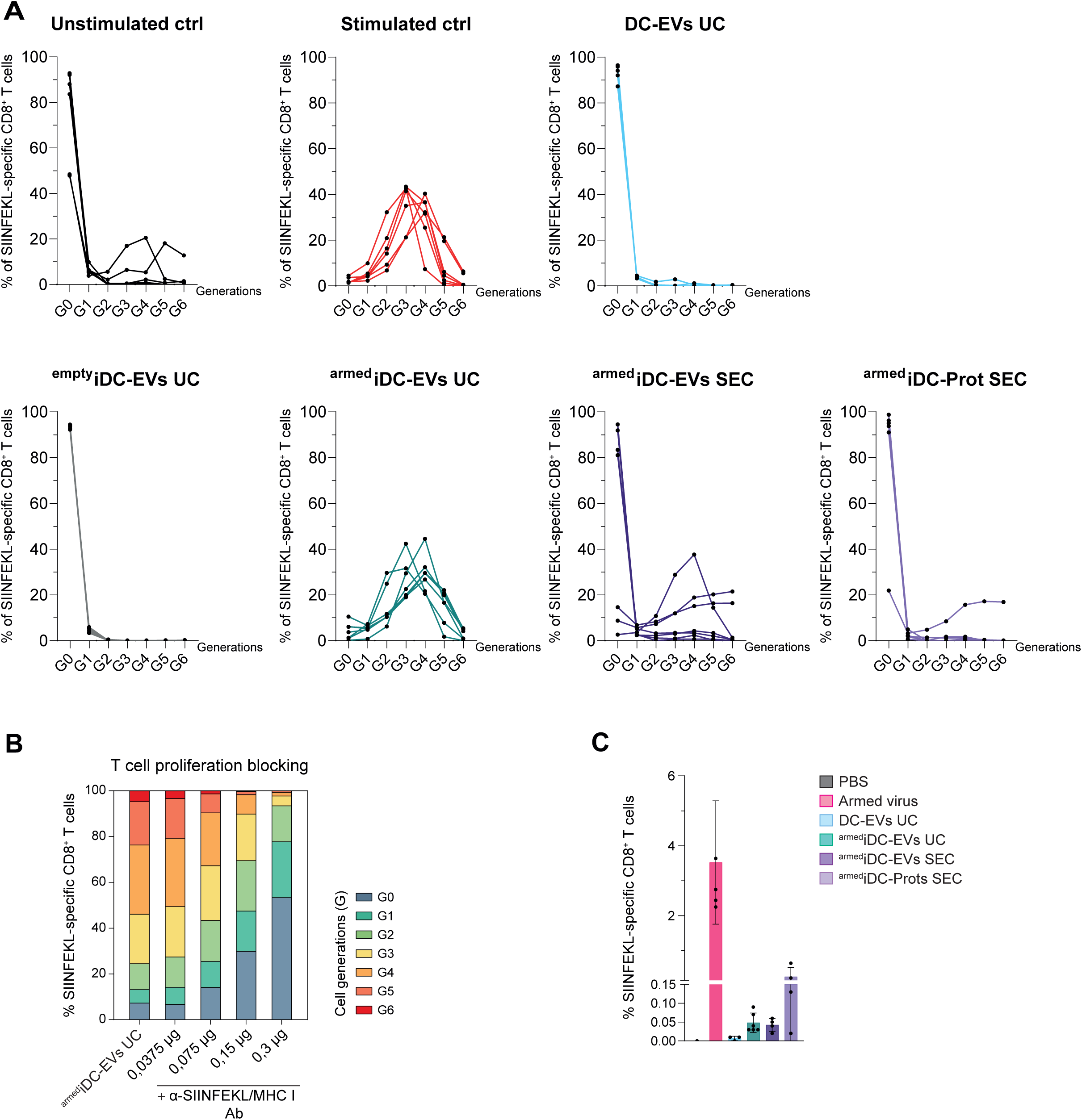
EVs isolated from therapeutic virus-infected dendritic cells activate and stimulate anti-tumoral cytotoxic T cells *in vitro*. A) Quantification of the number of CD8^+^ T cells per generation (from G0 to G6) following stimulation with positive control (anti-CD3/CD28 antibodies) or with EVs. One curve represents one experiment. B) CD8^+^ OVA-specific T cells proliferation by EVs relies on SIINFEKL presentation by MHC-I. T cells were incubated with EVs isolated by UC from DC2.4 cells infected with a triple armed MVA (^armed^iDC-EVs) and with increasing concentrations of anti-SIINFEKL/MHCI blocking antibody. Mean of n=2 values per condition. C) Cytometry quantification of the proportion of OVA-specific CD8^+^ T cells isolated from the circulating blood of tumor-bearing mice treated with a triple armed MVA virus or with EVs isolated from non-infected DC2.4 cells (DC-EVs UC) by UC, or from DC2.4 cells infected with a triple armed MVA by UC (^armed^iDC-EVs) or by SEC (^armed^iDC-EVs SEC for EVs and ^armed^iDC-Prots SEC for the soluble fraction). Blood was collected at day 21 from mice described in Figure 5. n=1 to 6 mice per group, in N=1 experiment.

